# Regulation of protein secretion through chemical regulation of endoplasmic reticulum retention signal cleavage

**DOI:** 10.1101/2021.10.19.464966

**Authors:** Arne Praznik, Tina Fink, Nik Franko, Jan Lonzarić, Mojca Benčina, Nina Jerala, Tjaša Plaper, Samo Roškar, Roman Jerala

## Abstract

Secreted proteins, such as hormones or cytokines, are key mediators in multicellular organisms. Protein secretion based on transcriptional control is rather slow, as proteins requires transcription, translation, followed by the transport from the endoplasmic reticulum (ER) through the conventional protein secretion (CPS) pathway towards the plasma membrane. An alternative faster bypass would be valuable. Here we present two genetically encoded orthogonal secretion systems, which rely on the retention of pre-synthesized proteins on the ER membrane (membER, released by cytosolic protease) or inside the ER lumen (lumER, released by ER luminal protease), respectively, and their release by the chemical signal-regulated proteolytic removal of an ER-retention signal, without triggering ER stress due to protein aggregates. Design of orthogonal chemically-regulated split proteases enables precise combination of signals into logic functions and was demonstrated on a chemically regulated insulin secretion. Regulation of ER escape represents a platform for the design of fast responsive and tightly-controlled modular and scalable protein secretion system.

Abstract figure:
membER and lumER system. By equipping a protein of interest (POI) with an N-terminal signaling sequence, which initiates the transport of proteins into the endoplasmic reticulum (ER), and a C-terminal KDEL ER retention sequence for luminal proteins or a KKXX sequence for transmembrane proteins, we can retain those proteins inside the ER and cis-Golgi apparatus (GA) through retrograde transport. Insertion of a protease cleavage site adjacent to the retention signal allows for the regulated fast secretion through proteolytic cleavage. The membrane bound, ER membrane (membER) and ER-luminal (lumER) systems allow for the controlled secretion of pre-synthesized protein, stored inside the ER. This platform enables release of target proteins several hours faster than systems relying transcription and translation.

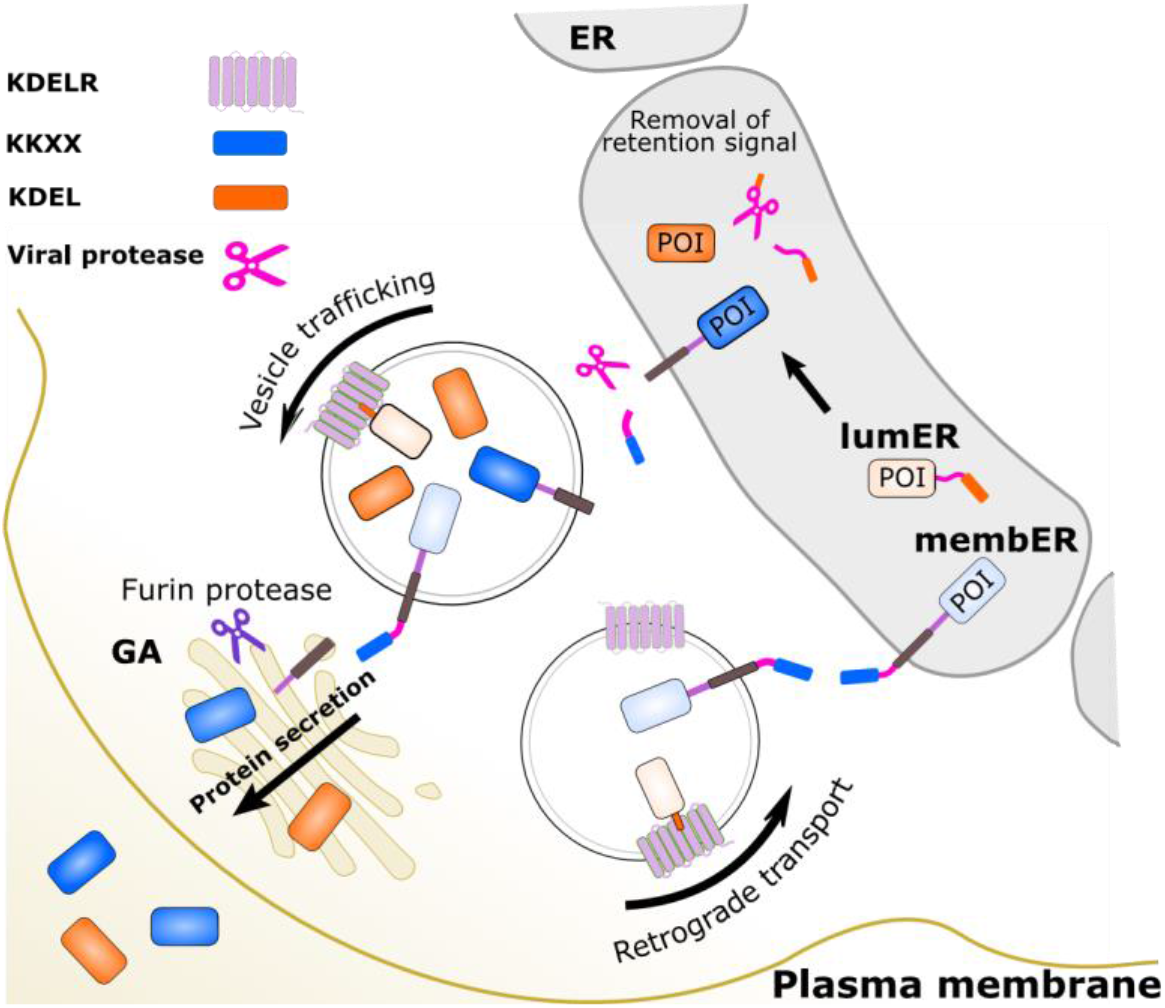

## Introduction

Protein secretion is an important cellular mechanism to translocate soluble or membrane bound proteins to respond to the changes in the cellular environment or signals from other cells, thus influencing a wide range of biological functions. Indeed, it has been estimated, that out of the ~ 25,000 full length open-reading frames (ORF) on the human genome, ~ 11% code for secretory proteins, with a further 20% coding for transmembrane proteins^1^. Regulated secretion of a protein of interest (POI) would offer an attractive tool for a diverse range of functions, such as hormonal release, homeostasis, immune defense, cytokinesis^2^ and quorum sensing^3^, as well as advanced therapeutic applications^4,5^. Several attempts have been made to control protein secretion, mostly by utilizing the established systems which rely on the control of gene expression^6–9^. These systems, while effective at regulating the production of proteins, are rather slow since they regulate gene expression rather than protein trafficking.

The processes of transcription and translation initiated by these systems can require several hours before the POI is secreted to a functionally relevant concentration. Natural processes, which need to respond rapidly to changes in the environment, often rely on the secretion of preformed proteins, stored in intracellular compartments and secreted quickly from the specialized cells by the regulated trafficking systems. Examples of such systems include the release of neurotransmitters into the synaptic cleft^10^ and insulin secretion from β-cells in the pancreas triggered by the increased concentration of glucose^11^. While the aforementioned cells contain specialized granules and machinery, capable of exporting proteins in response to a defined signal, most, if not all, eukaryotic cells possess the ability to secrete proteins from the cell through natural secretion pathways. The best characterized of these is the conventional protein secretion (CPS) pathway, where proteins pass through the endoplasmic reticulum, towards the Golgi apparatus (GA), the trans-GA and subsequently to the plasma membrane^2^. Early in their synthesis, an N-terminal or, alternatively, an internal signaling sequence on the newly forming protein marks the protein to be directed towards the ER lumen, where it passes through to GA in Coat-protein complex II (COPII)–coated vesicles^12,13^.

However, not all proteins that enter the ER are secreted – some proteins exert their function inside the ER and thus need to be retained there or recycled between the ER and the post-ER compartments. These proteins are typically marked with specific peptide tags, which distinguishes them from the proteins destined for secretion. The retention signal is recognized by specific receptors present in the ER and cis-GA, which retrieve the protein from post-ER compartments and return them to the ER by a retrograde translocation in COPI-coated vesicles. Examples of such sequences in animals include the C-terminal sequence motif Lys-Asp-Glu-Leu (KDEL), which is recognized by the KDEL receptor (KDELR) and is present on soluble secretory proteins, and the Lys-Lys-X-X sequence (KKXX, where X stands for any amino acid), which interacts with the recycling COPI protein complex and is often present at the cytosolic side of transmembrane proteins^14^. In principle, marking a POI with an N-terminal signaling sequence and the C-terminal retention signal could therefore load and localize a POI inside the ER.

Removal of the retention signal could provide an alternative system for controlling protein secretion. We reasoned that an important advantage of such a system would be faster response than that achieved by the induction of gene expression, relying instead on the pre-synthesized protein pool stored in the lumen of the ER. Proteolytic removal of the ER retention signal by a regulated protease could therefore transduce the signal into protein secretion. Several viral proteases from the *Potyviridae* family have been recently described and utilized as functionally inactive split proteins, which regain their catalytic activity upon reconstitution, triggered by several signals^15^. Since the viral proteases have a highly specific target sequence, they do not interfere with other cellular processes and are therefore valuable tools in synthetic biology for the generation of artificial protein-processing systems based on post-translational modifications^15,16^. Here, we developed a system for the controlled release of pre-synthesized proteins localized in the ER-lumen of mammalian cells, by proteolytic removal of an ER-retention signal sequence, based on the chemically regulated reconstitution of split potyviral proteases. Such a system is completely genetically encoded and its activation is considerably faster than systems based on the transcriptional induction of protein synthesis, since it relies on the pre-synthesized pool of proteins.

## Results

### Design of membER and lumER secretion system

Two types of ER-localized and proteolysis-based protein secretion systems were designed, each featuring a different C-terminal retention signal; the first type, termed membER, was designed for membrane-bound proteins retained in the ER by the addition of a transmembrane domain and a C-terminal KKXX sequence exposed to the cytosol (Figure 1a). This system is modelled after naturally occurring membrane-bound proteins localized in the ER. Based on previous reports, we opted to use the Lys-Lys-Met-Pro (KKMP) and Lys-Lys-Tyr-Leu (KKYL) sequences in the membER system, which were shown to be particularly potent Golgi-to-ER retrieval signals^17^. A transmembrane domain, from the B-cell antigen receptor complex-associated protein beta chain (CD79B) was used to anchor the protein of interest to the ER membrane. A potyviral protease cleavage site was inserted between the transmembrane domain and the ER retention signal, allowing for the control of secretion by the removal of the retention signal through proteolytic cleavage by cytosolic protease.

**Figure 1:**
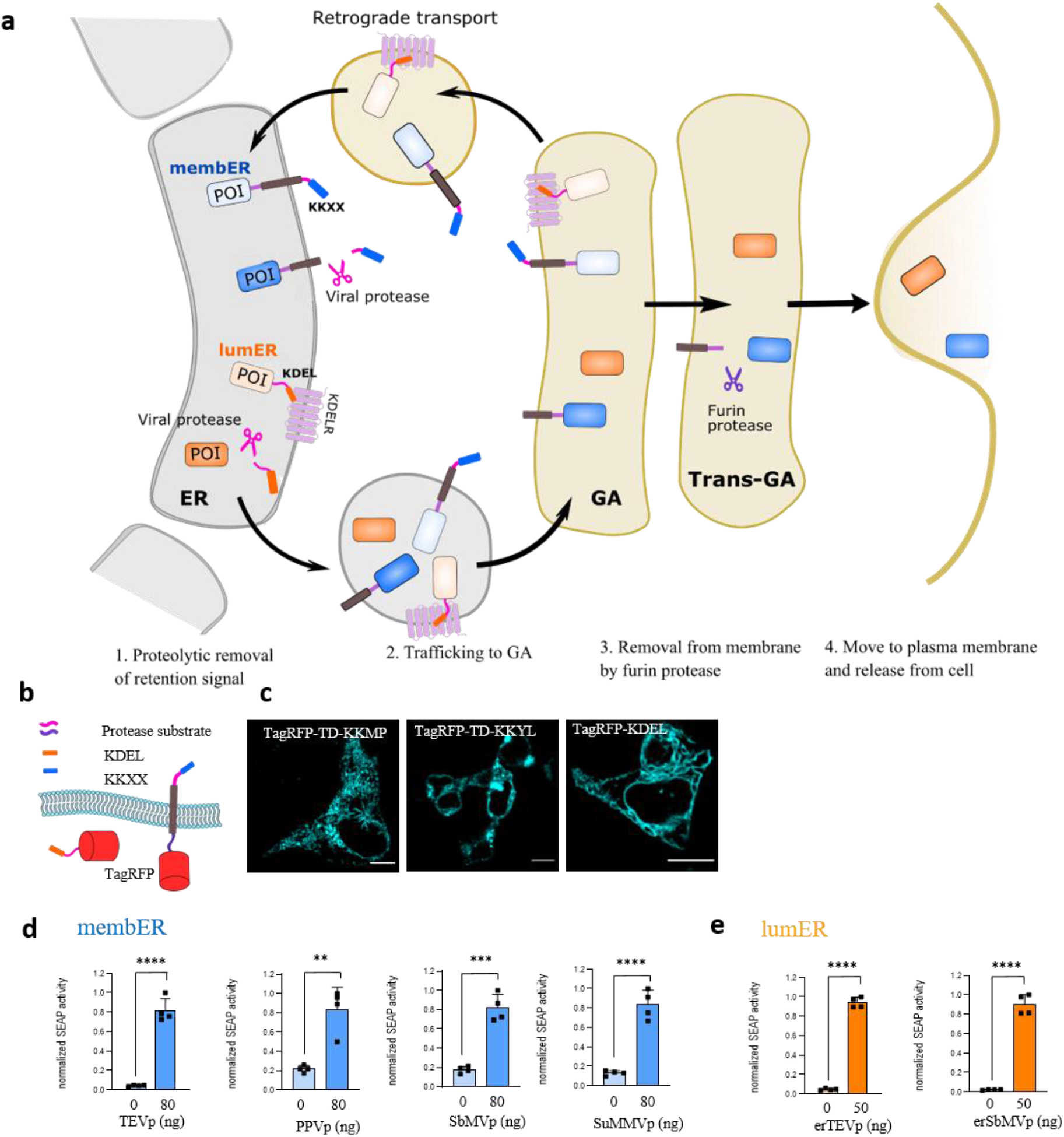
Protease-inducible protein secretion by the membER and lumER systems. **a**, Protease-triggered secretion of pre-synthesized protein from the ER is achieved by the cleavage of the C-terminal ER retention signal. A specific and orthogonal potyviral protease cleavage site was inserted between the protein of interest (POI) and the ER retention signal. Secretion is induced by the cytosolic protease (KKXX, member system) or ER-localized protease (KDEL, lumER system). A furin protease (FURp) cleavage site was inserted between the reporter and transmembrane domain to release the POI from the membrane for the member system. **b**, Design of the membER and lumER constructs featuring a TagRFP protein, which were used in microscopy imaging. **c**, Imaging of TagRFP, featuring a transmembrane domain and a C-terminal KKMP, KKYL or KDEL sequence. **d**, SEAP activity in cell supernatant was measured from cells transfected with membER constructs SEAP-FURs-TM-TEVs-KKYL, SEAP-FURs-TM-PPVs-KKYL, SEAP-FURs-TM-SbMVs-KKYL or SEAP-FURs-TM-SuMMVs-KKYL and co-transfected with TEVp, PPVp, SbMVp or SuMMVp, respectively. **e**, SEAP activity in the cell media from cells transfected with lumER constructs SEAP-TEVs-KDEL or SEAP-SbMVs-KDEL was measured after the co-transfection of an ER-localized TEVp (erTEVp) or erSbMVp, respectively. Values are the mean of four cell cultures ± s.d. An unpaired two-tailed t test (after equal variance was assessed with the F test assuming normal data distribution) was used for the statistical comparison of the data. Results are representative of two independent experiments.

The second type of an inducible protein secretion system, termed lumER, was designed for soluble proteins retained inside the ER lumen through the addition of a KDEL retention signal at their C-terminus (Figure 1a). Binding of the KDEL sequence by the KDELR (and thus recycling back to the ER) is pH sensitive, with maximal binding at pH below 6^18^. The acidic pH 6.2 inside the cis-Golgi lumen initiates binding, while the more neutral pH 7.2 to 7.4 inside ER lumen^19,20^ releases the interaction between KDEL and KDELR. KDELR itself features a KKXX retention signal at its cytosol-facing C-terminus, which interacts with the COPI vesicle machinery, responsible for the retro-transport of COPI vesicles from cis-Golgi to the ER^21^.

Both membER and lumER constructs were equipped with a signal peptide from the human immunoglobulin kappa^22^ at the N-terminus of the secretion constructs to ensure translocation of the protein into the ER during translation (sequences in Supplementary table 1-3). However, because the two retention systems work through different pathways, each of them required additional unique design considerations. For the membER system, removal of the retention signal enables the membrane-bound protein to bypass the cis-Golgi retro-transport, but once the protein passes through the GA, the protein remains anchored into the cell membrane due to the transmembrane domain. To achieve the release of the POI from the plasma membrane once it had passed the GA and before reaching the plasma membrane, an additional furin protease (FURp) cleavage site (amino acid motif R-X-X-R^23^) was inserted in the ER lumen-facing side of the target construct between the POI and the transmembrane domain. Since FURp is enriched in the GA^24^, it cleaves the POI off the membrane, enabling its release upon the fusion of the secretory vesicles with the plasma membrane^25–29^. The proteins presenting solvent-exposed R-X-X-R motif would be therefore cleaved on the way towards the plasma membrane.

On the other hand, the lumER secretion design relies on retention in the ER by the KDELR, which recognizes and binds the ER lumen-exposed C-terminal KDEL signal. For this reason, the lumER constructs do not need to be anchored to the ER-membrane to be retained and thus only require the N-terminal signal peptide and the C-terminal KDEL sequence, separated from the POI by a selected (chemically regulated) protease cleavage site (Abstract figure, Figure 1a). However, the proteolytic processing in the lumER system occurs inside the ER lumen, thus requiring a protease, which retains its location and activity within the secretory pathway. Additionally, this protease would need to be equipped with a signal sequence, for the transport into the ER, and a retention signal to halt its progression to the cell surface. We found a previously described ER-localized TEVp, termed erTEVp, appropriate for execution of the proteolytic processing inside the ER lumen^30^ that could be regulated by a chemical signal. To expand the repertoire of functional proteases used in our secretion systems, we replaced the TEVs in the membER systems with cleavage sites of other orthogonal viral proteases from the *Potyviridae* family of viruses, namely plum pox virus protease (PPVp)^31^, soybean mosaic virus protease (SbMVp)^32^, and sunflower mild mosaic virus protease (SuMMVp)^33^, which have been previously utilized in SPOC, the synthetic protease and coiled-coil-based signal processing system^15^. In a similar fashion to the described erTEVp, an ER-localized SbMVp and PPVp were designed to also feature an N-terminal signal sequence and C-terminal KDEL sequence for use with the lumER system.

### Protease-dependent release of proteins from mammalian cells

For visualization of trafficking of the membER and lumER constructs within the cell, TagRFP^34^ was fused to secretion constructs and expressed in HEK293T cells. (Figure 1 b-e, Supplementary figure 1a). For detection of proteins released into the cell media upon co-expression of an appropriate protease, reporter proteins secreted alkaline phosphatase (SEAP) and Gaussia luciferase (Gluc) were incorporated into the lumER and membER systems. The expression of both lumER-SEAP and membER-SEAP were verified by Western blot (Supplementary figure 1b, c). Since the implementation of KKYL retention signal produced less leakage compared to the KKMP (Figure 1d, Supplementary figure 2a), we opted for the use of the former retention signal in the membER system. We found that all four tested potyviral proteases were capable of inducing secretion when co-transfected with a membER construct containing the appropriate substrate. However, TEVp provided the best fold increase of the secreted reporter into the media in the presence of the protease (Figure 1d). Since secretion of the membER system depends on FURp processing to release the protein from the plasma membrane, cells were cotransfected with a plasmid expressing the native FURp. A further increase in the amount of a secreted protein was observed when FURp was added, without increasing the protein secretion in the absence of a protease (Supplementary figure 2b). In summary, the implementation of the membER secretion system with 4 orthogonal and highly specific proteases was demonstrated.

LumER-based SEAP secretion is controlled by proteolytic processing by an ER-localized protease. Reporter secretion was minimal in the absence of a potyviral protease, while the addition of erTEVp led to the 20-fold release of SEAP into the media (Figure 1e). Similarly, we tested PPVp and SbMVp on the lumER system, by replacing the TEVs with PPVs and SbMVs, and directing the two proteases into the ER lumen through inclusion of a signal peptide and a KDEL sequence. Coexpression of ER-targeted PPVp however did not lead to the release of SEAP into the cell media (Supplementary figure 2c). A large proportion of proteins travelling through ER and GA is processed by a diverse array of post-translational modifications, most notably glycosylation by a variety of ER- and GA-resident glycosyltransferase enzymes, which can modify proteins^35^. Glycosylation might decrease or altogether abolish the activity of a protein, depending on its location. Indeed, TEVp required mutations at two sites to prevent glycosylation and maintain its activity inside the ER lumen^30^. An online N-glycosylation site prediction tool NetNGlyc 1.0 Server (http://www.cbs.dtu.dk/services/NetNGlyc/) revealed two potential glycosylation sites in PPVp (N23, N173), potentially explaining the observed lack of activity of this protease in the ER. After changing the two amino acid residues (N23Q, T173G) at positions corresponding to erTEVp, small increase of secretion was observed (Supplementary figure 2d). ER-targeted SbMVp, on the other hand, was able to induce strong secretion of SEAP (Figure 1e), demonstrating here for the first time the activity and retention of SbMVp in the ER, therefore providing an additional orthogonal regulator of protein secretion.

Increasing the amount of a TEVp in combination with the membER constructs led to the increased secretion of SEAP (Supplementary figure 2e). On the other hand, increasing the amount of the ER-localized protease in the lumER system above a certain threshold led to a decreased secretion of SEAP (Supplementary figure 2f). We reasoned that this might be due to the competitive binding of the erTEVp to the KDELR, which also features a KDEL retention sequence on its C-terminus, and could lower the capacity of the retrograde transport. This was indeed confirmed by rescuing reporter secretion by the ectopic expression of the KDELR1, which led to an almost 2-fold increase of the amount of secreted SEAP triggered by the erTEVp (Supplementary figure 2g).

### Kinetics of the membER and lumER secretion systems

We next aimed to regulate secretion using chemically inducible proteases. For this purpose, the FKBP/FRB and ABI/PYL1 heterodimerization domain pairs were sued as chemically induced dimerization (CID) systems regulated by rapamycin^36^ and abscisic acid (ABA),^37^ respectively. Both CID systems were used to reconstitute split fragments of the four aforementioned pottyviral proteases, which regained their activity upon heterodimerization (Figure 2a). For use in the ER lumen, we designed pairs of split protease fragments of erTEVp, a signal sequence and a KDEL retention signal, fused to CID domains. The resulting pairs were termed er-rapa-TEVp and er-aba-TEVp (Figure 2a). HEK293T cells expressing the membER secretion system with SEAP as the POI and rapa-TEV or aba-TEV as the inducible split protease, secreted SEAP when stimulated with rapamycin or ABA, respectively (Figure 2b). As with single chain proteases, the KKYL retention signal resulted in higher fold of induction than KKMP (Figure 2b, Supplementary figure 3).

**Figure 2:**
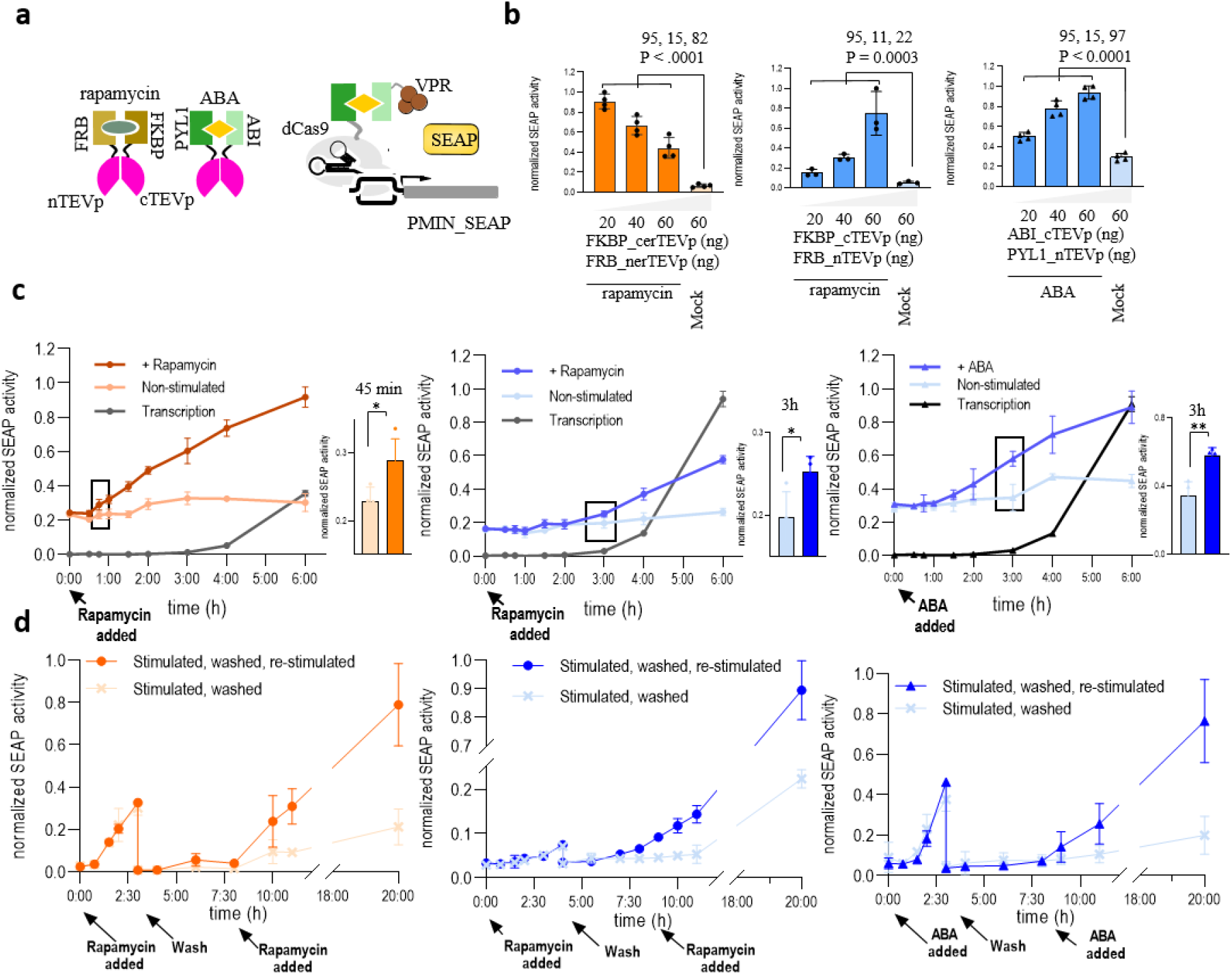
Inducibility and kinetics of protein secretion. **a**, For the inducible protein secretion, split protease fragments were fused to the chemically inducible dimerization partners FKBP/FRB or ABI/PYL1. For localization within the ER lumen, the inducible proteases were equipped with an N-terminal signal sequence and a C-terminal KDEL. Reconstitution of the protease was achieved by the addition of rapamycin in the case of FKBP/FRB or ABA in the case of ABI/PYL1. To compare the kinetics of secretion of our pre-synthesized proteins with secretion initiated by SEAP expression, a dCas9-FKBP/FRB-VPR or dCas9-ABI/PYL1-VPR system with an addition of an sgRNA targeting a minimal promotor upstream of the SEAP reporter gene, was utilized, where expression of SEAP was initiated by either rapamycin or ABA, respectively. **b**, Cells were transfected with SEAP-TEVs-KDEL and FKBP-cerTEVp/FRB-nerTEVp (er-rapa-TEVp; orange), SEAP-FURs-TM-TEVs-KKYL and FKBP-cTEVp/FRB-nTEVp or ABI-cTEV/PYL-nTEV (blue). SEAP activity was measured in the media after 20h after the addition of rapamycin or ABA. **c,** Secretion of SEAP was monitored after the addition of rapamycin or ABA for the KDEL (orange) or KKYL (blue) system and compared to the expression-driven secretion. **d**, Secretion was induced by the addition of rapamycin or ABA. After 4h, the medium was removed, the cells were washed and supplementary medium was added. After an additional 3h cells were restimulated. SEAP activity in the media was measured continuously. Values are the mean of four cell culture experiments ± s.d. Significance was tested by an unpaired two-tailed t test or one-way analysis of variance (ANOVA) with Tukey’s comparison (values of confidence intervals, degrees of freedom, F and P are indicated). Results are representative of two independent experiments.

Chemical stimulation of cells expressing er-rapa-TEV and the lumER secretion system resulted in an inducible secretion of SEAP. Similar to the single chain protease, higher amount of er-rapa-TEV resulted in a decreased amount of secreted SEAP after stimulation with rapamycin (Supplementary figure 4a), which can be attributed to the competition with the KDEL receptor. In case of er-aba-TEV heterodimerization domains within the ER, no secretion was detected for ABA stimulation (Supplementary figure 4b). The N-glycosylation site prediction tool indicated several potential glycosylation sites on both ABI and PYL1 heterodimerization domains (ABI: N292; PYL1: N66, N90), which likely hindered its function in the ER.

Rapamycin, on the other hand, was able to induce secretion already at picomolar concentrations and reached a plateau at approximately 25 nM with the membER system and 10 nM with the lumER system, while ABA was able to induce secretion in the membER system at nanomolar concentrations and reached a plateau at approximately 0.5 mM concentration (Supplementary figure 5). Taken together, both systems are able to elicit a concentration-dependent secretion of POI, however ABA only for the membER.

To monitor the secretion kinetics, cells harboring constructs for secretion systems and split chemically inducible TEVp were stimulated either with rapamycin or ABA and cell medium was harvested at designated time points. To directly compare the kinetics of a chemically-regulated secretion system with a chemically-regulated transcriptional activation system, we placed the expression of SEAP under a minimal promoter that was regulated by an inducible CRISPR-based transcription factor. A dCas9 protein fused to FKBP or ABI (dCas9:FKBP and dCas9:ABI) was brought into the proximity of the promoter by an sgRNA targeting the minimal promoter sequence. Transcription was initiated by the addition of rapamycin- or ABA-inducing heterodimerization of the Cas9 fusion with the transcription activator domain VPR fused to the complementary CID protein (FRB:VPR or PYL1:VPR, Figure 2a). Secretion kinetics for membER, lumER and the transcriptional activation-based systems were measured and compared at varying concentrations of the chemical inducers (Supplementary figure 6). While the proteolysis-based secretion systems were able to induce secretion in less than 1h, the transcription-activation based secretion was only observed after several hours. The lumER system was able to induce secretion with faster kinetics than the membER system; in the former a statistically significant increased amount of SEAP in the cell media compared to non-stimulated cells was observed at 45 minutes (Figure 2c). While the additional co-transfection of FURp with the membER system demonstrated increased secretion after extended time (Supplementary figure 7), it did not significantly increase the secretion kinetics in the first 4h (Supplementary figure 2b).

We next tested the reversibility of both secretion systems. After induction with either rapamycin or ABA, cell media were sampled for up to 3-4h, after which the cells were washed and fresh media were supplemented. SEAP concentrations continued to remain low in the media up until 4h after washing, after which cells were restimulated and a repeated secretion of SEAP was observed in stimulated cells, with minor leakage from the unstimulated cells even after 20h (Figure 2d), suggesting that both system could be activated as multiple pulses.

Next, we tested how long it takes to release the total amount of the ER-localized protein pool from the cell after the induction of secretion. Since the constitutive expression of our constructs is continuously replenishing the ER with the POI, we aimed at halting the transport of newly formed protein into the ER lumen, such that only the pre-synthesized protein pool could be secreted. Cells were treated with Eeyarestatin I (Sigma-Aldrich, E1286), a potent inhibitor of Sec61-mediated protein translocation into the endoplasmic reticulum, which has been shown to inhibit secretion of cellular proteins at the ER-resident ribosomal biosynthesis step^38^. Media were harvested after induction of secretion with rapamycin. We observed that the majority of the available SEAP was secreted within 6-7 h after stimulation for both membER and lumER system (Supplementary figure 8).

The intracellular trafficking of membER and lumER proteins within the cell was monitored under the fluorescence microscope, by the inclusion of a super-folder GFP (sfGFP), split into two segments between the 10^th^ (GFP1-10) and 11^th^ β-strand (GFP11), which have been previously shown to allow for a high resolution imaging within cells^39^. GFP1-10 was fused to the membER (SS-GFP1-10-FURs-TD-TEVs-KKYL) and lumER (SS-GFP1-10-TEVs-KDEL) secretion systems, while seven tandem repeats of GFP11 (GFP11×7) were fused to the human Fas receptor transmembrane domain. The trafficking of the ER-localized construct to GA and further to the plasma membrane could be visualized be the reconstitution of the split sfGFP when they were localized in the same cellular compartment (Supplementary figure 9a, 10a). Since the GFP11×7 fragment, once produced, needs to pass the ER to reach GA and the plasma membrane, the fluorescence could be already observable in the ER (Supplementary figure 9b, 10b). Both the lumER and membER systems retained the reconstituted protein within the ER when no protease was added. Co-expression of TEVp and erTEVp led to the appearance of a fluorescent signal at the plasma membrane (Supplementary figure 9c, 10c). The reconstitution of a split TEVp and split erTEVp similarly led to the accumulation of the split GFP fragments at the plasma membrane after 3h (Supplementary figure 9d, 10d). When visualizing a group of cells, a noticeable increase in fluorescence was observed in the GA in less than 15 minutes after stimulation with rapamycin, while the signal at the plasma membrane was observed after 1h (Figure 3).

**Figure 3:**
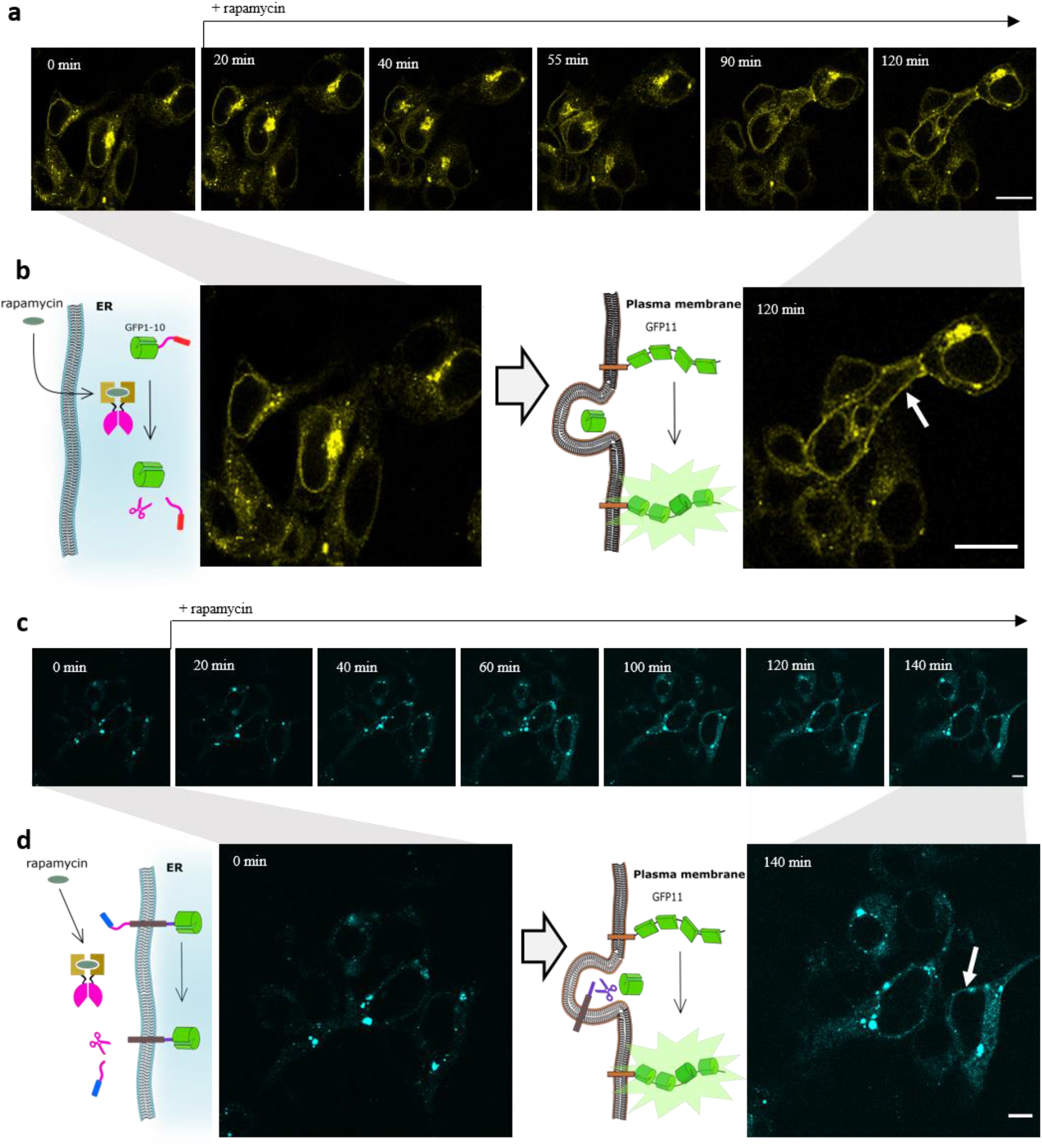
Single cell visualization of protein trafficking in the lumER and membER system. **a**, GFP signal was visualized on the same group of cells over a period of 2h. **b**, Close up of cells before the addition of rapamycin and of 2h after the addition of rapamycin. The signal begins to appear at the plasma membrane (white arrow) after ~ 60 min. **c,** GFP signal was visualized on the same group of cells over a period of 140 min. The signal begins to appear at the plasma membrane (white arrow) after ~ 110 min. **d**, Close up of cells before the addition of rapamycin and of cells 140min after the addition of rapamycin with an arrow indicating localization at plasma membrane. Scale bar = 10 μm

Since the two systems are based on differently localized proteases and different recycling signals it is likely that they could be regulated independently. Orthogonality of the two secretion systems was tested by co-transfecting both the lumER and membER systems simultaneously. For the membER system a SEAP reporter was used, while the lumER was fused to a Gluc reporter (Figure 4a). Cells expressing both systems were cotransfected with TEVp, erTEVp or both proteases and measured Gluc and SEAP activity in the media. Each system elicited secretion only in the presence of the appropriate protease (Figure 4b). Similarly, we managed to elicit the selected secretion by co-transfecting a cytosolic ABI-cTEVp/PYL1-nTEVp and ER localized er-rapa-TEVp (Figure 4c) and inducing secretion by the addition of either ABA or rapamycin (Figure 4d).

**Figure 4:**
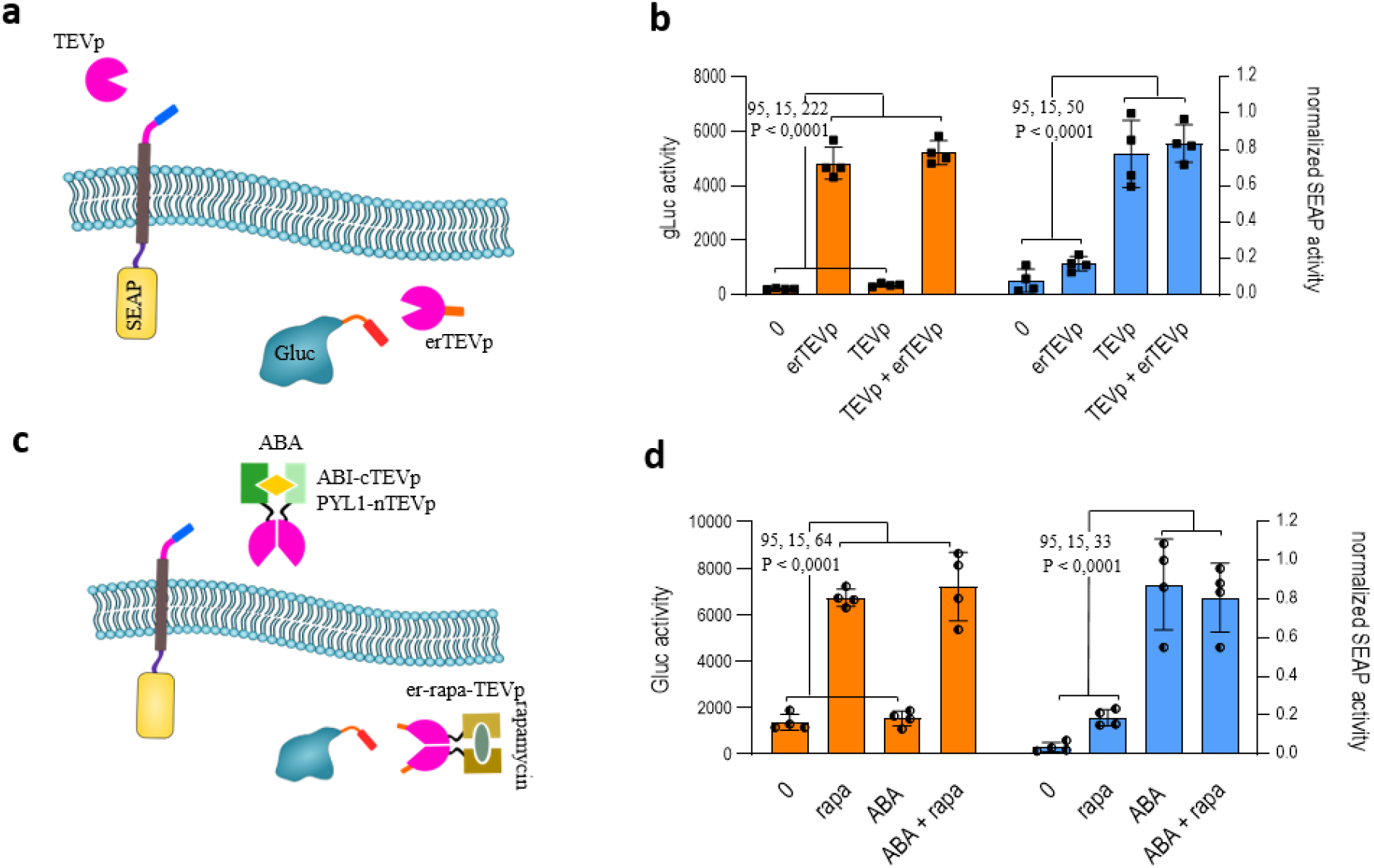
Orthogonality of the lumER and membER system. **a**, Cells were co-transfected with membER system, featuring SEAP as a reporter, and lumER system, featuring a Gluc reporter. **b**, Gluc (orange) and SEAP (blue) activity was measured in the media after the addition of TEVp and erTEVp. **c**, Design of the inducible and orthogonal system. ABI-cTEVp/PYL1-nTEV and er-rapa-TEVp were co-transfected with the membER-SEAP and lumER-Gluc constructs. **d**, the systems were tested with inducible proteases. ABI-cTEV/PYL1-nTEV was used for secretion with the membER (blue) and FKBP-cerTEV/FRB-nerTEV was used for the lumER system (orange). Values are the mean of four cell cultures ± s.d and are representative of two independent experiments. Significance was tested by oneway analysis of variance (ANOVA) with Tukey’s comparison (values of confidence intervals, degrees of freedom, F and P are indicated).

### Construction of an input signal-processing module to regulate protein secretion via the membER and lumER systems

The ability to regulate the removal of the retention sequence with several chemical or biological signals would allow for a stringent control over protein secretion, as well as allow the system to be controllable by different combinations of external but also internal signals. While KKYL or KDEL is required at the C-terminus of the protein to be retained within the ER, only a single retention signal can be present on each protein, making it difficult to design a construct responding to a combination of two inputs. We therefore employed the previously described SPOC system^15^, based on split proteases and combinations of coiled-coil peptides, which is able to process two input signals according to Boolean logic operations and produce an output in the form of a reconstituted viral protease activity^15^. The reconstituted viral protease could therefore be further used to regulate protein secretion. The principle of SPOC logical circuits used in this study is described in the Supplementary figure 11. We employed the constructs to demonstrated an A nimply B function, activated by the presence of one signal and the absence of the other one (Figure 5a), as well as an AND function, activated by the presence of both signals using membER secretion system (Figure 5b), both with excellent difference between the active and inactive output state. However, any two- or multiple input Boolean logic could be implemented in principle.

**Figure 5:**
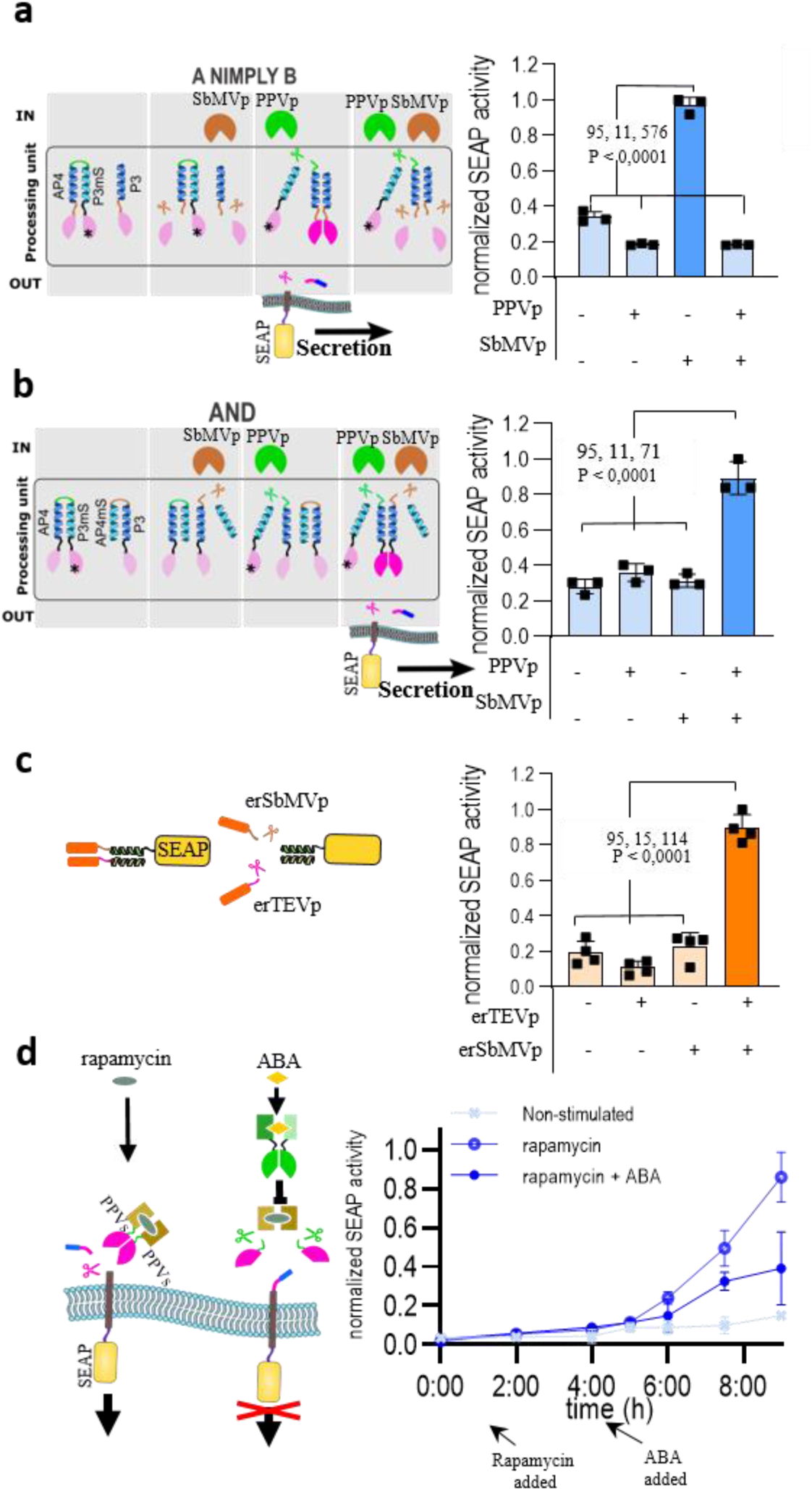
Modular combinations and two-input signal processing capacity of protein secretion systems. **a**, A two-input Boolean logic processing system - SPOC, previously described by Fink *et al*., was used to control the reconstitution of a split TEVp and regulate secretion through the removal of KKYL. The input proteases SbMVp and PPVp produced the output according to A nimply B logical function and **b**, AND logical function. For the KDEL system, SEAP was retained inside the ER by the addition of a C-terminal KDEL sequence, as well as the retention of the construct through the interaction of the P3/P4 coiled coils, where P3 also featured SbMVs and a KDEL sequence. Secretion was induced by the addition of erTEVp and erSbMVp. **c**, to establish an AND function for the lumER system, P3-TEVs-KDEL was co-transfected and localized inside the ER with SEAP-P4-SbMVs-KDEL. Apart from the KDEL sequence present on the same protein, SEAP was retained inside of the ER through the P3/P4 coiled-coil interaction. To facilitate the secretion of SEAP, both SbMVp and TEVp have to present, in order to remove the two retention signals. **d**, Implementation of a protein secretion OFF switch. Rapamycin is used to reconstitute a split TEVp, which further drives secretion of the membER secretion construct. By inserting a PPVp cleavage site (PPVs) between the dimerization domain and protease fragment, the reconstitution of TEVp can be inhibited with an ABA-inducible split PPVp. Cells were stimulated with rapamycin to induce secretion. After 3h ABA was added and cell media were harvested continuously for 6h. Values are the mean of four cell cultures ± s.d and are representative of two independent experiments. Significance was tested by one-way analysis of variance (ANOVA) with Tukey’s comparison (values of confidence intervals, degrees of freedom, F and P are indicated).

Optimization of each protease to retain the activity in the ER for lumER system could likely be accomplished by introduction of appropriate mutations in the proteases. More importantly, transferring the components of SPOC into the ER might increase leakage of the system due to the saturation of the recycling machinery. We therefore employed an alternative strategy to adapt SPOC to lumER and designed a system that allows retention of proteins in the ER and their release upon signal processing of two input signals. We appended the KDEL sequence to an ER-localized designed coiled-coil forming P3 peptide via a TEVp cleavage site (P3-TEVs-KDEL), which interacts with its complementary peptide P4, located on our POI (SEAP-P4). The interaction of P3/P4 coiled coil pair was capable of retaining SEAP within the ER when the KDEL sequence was present only on one of the two interacting constructs (Supplementary figure 12a). In this system, cleavage with TEVp resulted in release of a POI. To perform an AND function in a lumER system, an additional KDEL sequence was fused to the protein of interest with SbMVs inserted between P4 and KDEL (SEAP-P4-SbMVs-KDEL) (Supplementary figure 12b). This resulted in induction of secretion only when both erTEVp and erSbMVp are present (Figure 5c).

To further control secretion, we aimed at establishing a simplified version of an A nimply B logic function to function as an OFF switch, which was intended to halt secretion, at some selected delay after it had been initiated. For this purpose, we inserted a PPVs between the heterodimerization domains and the split TEVp fragments (FKBP_PPVs_cTEVs, FRB_PPVs_nTEVp). The addition of PPVp cleaves off the split protease fragments from the heterodimerization domains, preventing their reconstitution and, consequently, secretion (Supplementary figure 13 a, b). To make the OFF switch inducible, ABA CID heterodimerization domains were used to control the assembly of PPVp (ABI_cPPVp/PYL1_nPPVp) (Figure 5d). Cells which had been stimulated with rapamycin and were subsequently stimulated with ABA, demonstrated a decreased secretion of SEAP into the media compared to cells only stimulated with rapamycin (Figure 5d).

### Control of insulin secretion with the lumER system

Finally, we applied our secretion system on a therapeutically relevant protein, which could be used to induce fast release upon stimulation. Diabetes mellitus affects millions of people worldwide and is characterized by a depletion or exhaustion of insulin producing β-cells in the pancreas^40^. Insulin is normally secreted from specialized granules inside of β-cells and is tuned to be released within minutes to hours after the rise of blood glucose level^41^. Engineered systems have been previously designed to trigger the release of insulin or GLP-1, which however exhibited response time of several hours^6^.

We used a design of human insulin suited for protease processing and secretion through the conventional secretion pathway^42^. This was achieved by modification of the insulin sequence to feature two FURs surrounding the C-peptide, which could be processed to mature insulin by FURp. The preproinsulin sequence was appended to the lumER secretion system (Figure 6a) in combination with the er-rapa-TEV. Human insulin levels secreted by HEK293T cells were quantified by measuring the amount of secreted C-peptide upon rapamycin stimulation. Alternatively, a reporter system was designed, which substitutes the C-peptide of a preproinsulin with a short secretory Gaussia luciferase, and has been previously used as a surrogate reporter of insulin secretion^43,44^ (Supplementary figure 14). Elevated C-peptide levels were detected within cell media as early as 30 minutes after the addition of rapamycin (Figure 6b). The presence of a C-peptide in the cell media with the lumER system was detected within 45 minutes, in contrast to several hours before a system which relied on the transcriptional induction of insulin expression by the addition of rapamycin (Figure 6b).

**Figure 6:**
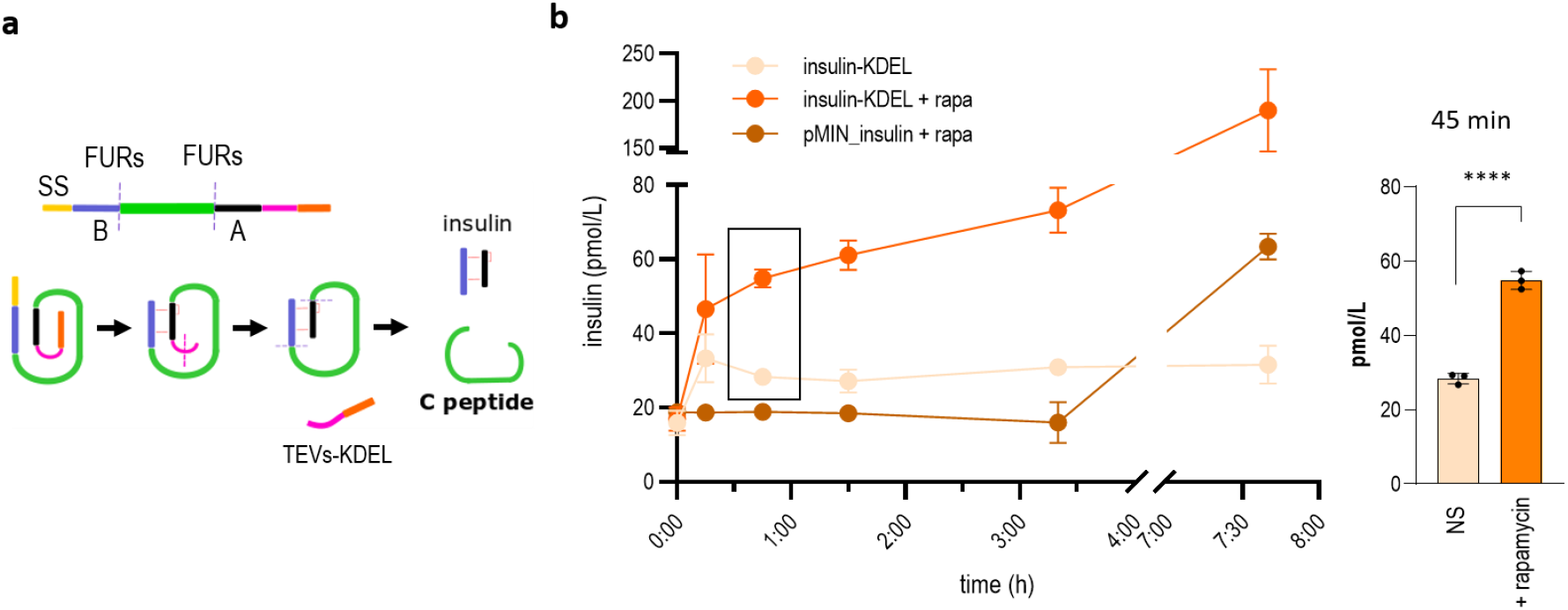
Inducible secretion of insulin based on the chemically regulated lumER system. **a**, A modified preproinsulin sequence was designed to feature FURs (purple lines) between B (blue) and C-peptide (green) and C and A-peptide (magenta). After removal of the retention signal, the proinsulin passes GA, where it is processed to the mature insulin by the FURp. Media were harvested at specified time points and C-peptide ELISA was performed to measure the amount of processed insulin. **b**, C-peptide concentration was measured in the cell media after the addition of rapamycin, as well as from non-stimulated (NS) cells. Secretion kinetics of the lumER-insulin system was compared to the kinetics of secreted insulin in an inducible transcription-activation system. The secretion was compared with cells, which were not stimulated with rapamycin. Values are the mean of three cell cultures ± s.d. and are representative of two independent experiments. An unpaired two-tailed t test (after equal variance was assessed with the F test assuming normal data distribution) was used for the statistical comparison of the data.

## Discussion

Several biological processes require response within minutes to hours, in order to elicit a physiologically relevant response, which could not be achieved using a transcriptional control. Examples of processes that require fast response include the release of other hormones depending on the physiological conditions, such as insulin after a meal, release of pro- and anti-inflammatory cytokines, anti-microbial or anti-viral peptides^45–51^. Therefore the development of faster protein secretion systems offers an attractive venue for delivering biotherapeutic agents *in situ* in a physiologicaly relevant timeframe^4^. Natural protein secretion systems, that are able to respond rapidly to input signals, often achieve this in specialized cells, by forming storage granules, containing pre-synthesized protein, which are positioned close to the plasma membrane and fuse with the plasma membrane upon signal detection. The mechanism of biosynthesis of secretory granules in specialized cells and the synthesis of their release components involves tens of genes and is still a relatively unknown process^52^ and those specialized cells may be triggered by external signals^53^. However, accelerated kinetics of selected protein release in non-specialized cells based on the principle of storing pre-synthesized proteins and releasing them upon stimulation is an attractive approach to generate synthetic system for protein release.

Here two orthogonal systems for the release of pre-synthesized proteins from the ER by the proteolytic removal of a C-terminal ER-retention signal were presented. Both the lumER and the membER system are able to secrete proteins substantially faster than engineered systems, which rely on the transcriptional regulation of secretory proteins. Both systems described here offer a tunable, concentration dependent release of proteins, which achieves the rate of secretion kinetics suitable for several faster-acting biological processes. We demonstrated the use of a chemically regulated TEVp inside the ER, as well as the activity of an additional highly specific and orthogonal viral protease, SbMVp, in the ER lumen. The lumER system proved to be the faster of the two, leading to protein secretion approximately 45 minutes after stimulation. This difference could be due to the requirement for processing by a single protease in lumER, whereas the membER system requires additional processing by the FURp. Additionally, the substrate and split protease concentration might be higher in the ER, compared to the cytosolic protease. The CPS trafficking kinetics and capacity itself poses a limit on the speed of secretion as proteins can be transported inside the transport vesicles from ER to GA and further on to the plasma membrane at a certain maximal rate^54^, preventing the system to reach the kinetics of granule release in specialized cells.

An important feature of this platform is that it relies on the removal of the retention signal by well defined, orthogonal and inducible proteases with high selectivity. This allows the systems to be incorporated into a variety of processing modules, which result in the activation or inactivation of an effector protease, thus allowing for precise control of protein secretion. As shown here, this processing can incorporate Boolean logic functions, and could potentially be expanded to feature other protein circuits^16^, feedback loops etc. Furthermore, the inducibility of the input protease could be controlled by a variety of heterodimerization domains^55^. This endows our system with a versatility not found in similar systems, which also rely on the accumulation of pre-synthesized proteins inside of ER, but retain the protein inside the ER through protein aggregation^29^. Importantly the system described here rely on monomeric proteins, avoiding protein aggregates that are known to trigger ER stress, which may lead to cell apoptosis^56^.

Post-translational modifications of proteins, which occur in the CPS pathway may interfere with designed proteins and certain systems need to be optimized when translocated to the ER, as demonstrated for the PPVp for the lumER system. While certain modifications can restore protein activity (as was the case with erTEVp^30^), not all proteins can be readily transformed to function within the ER, presenting a challenge which needs to be addressed individually for each system. However, we presented several functional modules such as the SbMVp and designed coiled-coil heterodimerization domains that could be combined into new functional circuits within the ER. On the other hand, certain post-translational processes can be used to the advantage of a designed system, such as the proteolytic cleavage by the GA-localized FURp, which we utilized for the release of proteins from the plasma membrane, in the case of the membER system, as well as for processing of the engineered proinsulin into insulin.

In summary, the presented platform of protein secretion regulation based on modulation of protein recycling to the ER through cleavage of the retention signal present several advantages as a modular and scalable system that could be used in mammalian cells and likely in eukaryotic cells in general.

## Materials and Methods

### Plasmid construction

Plasmids were constructed using the standard procedures of molecular cloning or Gibson assembly. The amino acid sequences of all constructs are provided in Supplementary table 1. All the protease coding sequences were codon optimized for expression in human cells, and the DNA was synthesized by IDT (PPVp, SbMVp, SuMMVp) or Life Technologies (TEVp). Split N- and C-fragments of proteases in fusion with CCs or dimerization partners FKBP/FRB and ABI/PYL1 were PCR amplified and inserted into the pcDNA3 vector. The FKBP and FRB domains were obtained from the pC4-RHE and pC4EN-F1 plasmids. ABI and PYL1 were obtained from pSLQ2816 pPB, as follows: CAG-PYL1-VPR-p2A-GID1-ABI-WPRE PGK-GAI-tagBFP-SpdCas9 (Addgene: 84261)^36^. The GFP1-10 and GFP11×7 fragments were obtained from plasmids pcDNA3.1-GFP(1-10) (Addgene: 70219) and pACUH-GFP11×7-mCherry-α-tubulin (Addgene: 70218)^39^.

Mutations N23Q, C130S and N171T in the wild type TEVp were made in the design of erTEV. SEAP was obtained from pPIRON-SEAP-pA which was kindly provided by Martin Fussenegger (Institute of Biotechnology, Swiss Federal Institute of Technology, ETH Zürich). The constructs for logical processing used with the membER system were described by Fink *et al*^15^.

The nuclease-deficient Cas9 was obtained from pHR-SFFVdCas9-BFP-KRAB (dCas9; plasmid 46911, Addgene). The gRNA containing MS2 aptamer sequences (gRNA_2 ms2) was synthesized by LifeTechnologies, amplified with PCR and inserted into the plasmid pgRNA-humanized. All gRNA target sites were designed as 7 or 10 copies of the bnding sites, separated by hypervariable linkers and synthesized by Genewiz, similar as previously described by Lebar *et al* (2020) ^57^. dCas9 was fused with the heterodimerization domain FKBP or ABI. The VPR activator sequence was adapted from the Addgene plasmid 63801 and synthesized by Genewiz. The transcription activator was fused with either FRB or PYL heterodimerization domains, respectively, to allow for activation of transcription by the addition of either rapamycin or ABA to the cell media. Detailed information on the sequences used in this study are presented in Supplementary tables 1-3.

### Cell culture, SEAP detection and luciferase assay

The human embryonic kidney (HEK) 293T cell line was cultured in DMEM medium (Invitrogen) supplemented with 10% fetal bovine serum (FBS; BioWhittaker, Walkersville, MD, USA) at 37°C in a 5% CO_2_ environment. Following this, 2 x 10^4^ HEK293T cells per well were seeded in clear 96-well plates (CoStar, Corning). At a confluence of 50–70% cells were transfected with a mixture of DNA and PEI (8 μl PEI/1,000 ng DNA; stock concentration 0.324 mg/ml, pH 7.5). Detailed information about the amounts of plasmids in each experiment is provided in Supplementary table 4.

Rapamycin (Sigma-Aldrich) and abscisic acid (ABA, Sigma-Aldrich) were each dissolved in dimethyl sulfoxide (DMSO) at concentrations 1 mM and 50 mM, respectively. Forty-eight hours after the transfection, cells were stimulated with rapamycin and/or ABA at a final concentration of 1.5 μM and 50 μM, respectively. At different time points after induction, media was harvested, centrifuged at 1200x g for 5 min to remove any leftover floating cells and transferred to a clear 96-well plate for SEAP detection or a white 96-well plate (CoStar, Corning) for luciferase assay. The remaining cells were harvested and lysed with 30 μl of 1× passive lysis buffer (Promega).

To measure the time it takes for the complete secretion of the pre-synthesized and ER-localized protein, HEK293T cells (5 × 10^5^ to 7 × 10^5^) were seeded in six-well plates (Techno Plastic Products) and transfected the next day. 48h after transfection cell media was changed and Eeyarestatin I (E1286) was added to a final concentration of 8 μM. After 1h cells were treated with rapamycin at a final concentration of 1.5 μM and 10 μl of cell media was harvested at specified time points.

SEAP was detected in the media and in the lysate with an alkaline phosphatase detection medium – powder QUANTI-Blue™ (InvivoGen) as per the products instructions. Briefly, 200 μl of the resuspended QUANTI-Blue™ powder was added to 10 - 30 μl of the harvested media or the cell lysate and incubated at 37°C for 1-2h. The absorbance of the media was measured at 630nm with a Synergy Mx (BioTek) microplate reader. Normalized SEAP activity was calculated by dividing the set of values by the highest value in the experiment. In the secretion kinetics experiment the values for the membER, lumER and transcription systems were normalized separately. Firefly luciferase and Gaussia luciferase activity were measured using a dual luciferase assay (Promega) and an Orion II microplate reader (Berthold Technologies). Luciferase activity presented as relative light units (RLU) was calculated by dividing each samples Gaussia luciferase activity, measured in the cell media, by the constitutive Firefly luciferase activity determined in the lysed cells from the same sample. Normalized Gaussia luciferase activity was calculated by dividing the set of values by the highest value in the experiment.

### Imaging and *in situ* monitoring of protein trafficking

5 x 10^4^ HEK293-T cells per well were seeded in an 8-well chamber slide (ibidi). At a confluence of 50–70% cells were transfected with a mixture of DNA and PEI (8 μl PEI/1,000 ng DNA). Detailed information about the amounts of plasmids in each experiment is provided in Supplementary table 4. Forty-eight hours after transfection, HEPES was added to a final concentration of 20 μM. For experiments of monitoring inducible protein trafficking, cells were treated with rapamycin and/or ABA at a final concentration of 1.5 μM and 50 μM, respectively. Cells were imaged using a Leica TSC SP5 confocal microscope equipped with an HCX plan apo 963 (NA 1.4) oil-immersion objective. During imaging, the cells were kept at 37°C. For acquisition, Leica LAS AF software was used and for image processing, LAS AF and ImageJ software were used.

### Monitoring insulin secretion with C-peptide ELISA

Recombinant human preproinsulin was designed according to Hay and Docherty^26^ to allow for the secretion of the peptide through the conventional protein secretion pathway. The preproinsulin sequence featured a furin protease cleavage site between B:C and C:A peptides, to allow for the processing to insulin inside of the GA. A TEV protease cleavage site and KDEL retention signal was fused to the C-terminal end of the recombinant insulin. 10^5^ HEK293T cells per well were seeded in clear 24-well plates (CoStar, Corning). At a confluence of 50–70% cells were transfected with a mixture of DNA and PEI (8 μl PEI/1,000 ng DNA). Detailed information about the amounts of plasmids in each experiment is provided in Supplementary table 4. Forty-eight hours after the transfection, cell media was removed and supplemented with fresh media. After 1h, cells were stimulated with rapamycin at a final concentration of 1.5 μM and media was harvested at specific time points after stimulation. The media was centrifuged for 5 min at 1200 r.p.m. to remove any floating cells and 100 μl media was transferred to a clear 96-well plate. Human insulin levels secreted from HEK293T cells were quantified using an Ultrasensitive C-peptide ELISA kit (Mercodia, cat. no. 10-1141-01) and the concentration was calculated from the standard curve based on the supplied calibrators.

### Transcriptional regulation of protein secretion

For comparing the secretion kinetics of our post-translational modification-based system with secretion systems based on inducing gene expression, we cloned our reporter proteins SEAP and Gluc, as well as recombinant insulin under a minimal promoter, which featured 7 repeats of target binding sites [b]. Cells were seeded and transfected at a confluence of 50–70% cells with the plasmid, carrying the desired reporter protein, as well as plasmids coding for dCas9-FKBP, FRB-VPR and gRNA in the case of a rapamycin inducible system or dCas9-ABI, PYL-VPR and gRNA in the case of an ABA inducible system. Detailed information about the amounts of plasmids in each experiment is provided in Supplementary table 4. At determined time points, the media was harvested and the reporter activity was measured or protein concentration was determined accordingly.

The human preproinsulin, modified to feature FURs between B:C and C:A peptides, was cloned under a minimal promoter, which featured 10 repeats of target binding sites [ab]. pMin_insulin was used in combination with dCas9-FKBP and FRB-VPR to control the transcription of insulin by the addition of rapamycin.

### Immunoblotting

HEK293T cells (5 × 10^5^ to 7 × 10^5^) were seeded in six-well plates (Techno Plastic Products). The next day, at a confluence of 50–70%, the cells were transiently transfected with a mixture of DNA and PEI (8 μl PEI/1,000 ng DNA). Detailed information on the amounts of plasmids in each experiment is provided in Supplementary table 4. At 48 h after transfection, the cells were washed with 1 mL PBS and lysed in 100 μL of lysis buffer (40 mM Tris-HCl, pH 8.0, 4mM EDTA, 2% Triton X-100 and 274 mM NaCl) containing a cocktail of protease inhibitors (Roche). Cells were lysed for 20 min on ice and centrifuged for 15 min at 17,400 r.p.m. to remove cell debris. The total protein concentration in the supernatant was determined with BCA assays. Proteins from the supernatant were separated on 12% SDS–PAGE gels (120 V, 60 min) and transferred to a nitrocellulose membrane (350 mA, 90 min). Membrane blocking, antibody binding and membrane washing were performed with an iBind Flex Western device (Thermo Fisher) according to the manufacturer’s protocol. The primary antibodies were rabbit polyclonal anti–AU1 tag (Abcam ab3401; diluted 1:2,000). The secondary antibodies were HRP-conjugated goat anti–rabbit IgG (Abcam ab6721; diluted 1:3,000) and HRP-conjugated goat antimouse IgG (Santa Cruz, sc2005; diluted 1:3,000). The secondary antibodies were detected with ECL western blotting detection reagent (Super Signal West Femto; Thermo Fisher) according to the manufacturer’s protocol.

### Software and statistics

Graphs were prepared with GraphPad Prism 8 (http://www.graphpad.com/). GraphPad Prism 8 was also used for statistical purposes. Values are the means of at least three/four biological replicates ± standard deviation (s.d.) and are representative of at least two independent experiments. An unpaired two-tailed *t*-test (equal variance was assessed with the *F*-test assuming normal data distribution), one-way analysis of variance (ANOVA), Tukey’s comparison and Dunnett’s comparison were used for the statistical comparison of the data.

### Data availability

The authors declare that the data supporting the findings of this study are available in the paper and its supplementary information files. The raw data are available from the corresponding author upon reasonable request.

## Acknowledgements

The project was founded by Slovenian Research Agency (program P4-0176, projects J1-9173, Z4-2657, N4-0800) and European Research Council (ERC AdG MaCChines 787115). This work was initiated as part of the 2016 iGEM competition and we would like to thank the Slovenian 2016 iGEM team members who are not listed as authors – M. Meško, T. Lebar, F. Lapenta, Ž.Strmšek, R. Krese, K. Leben, L. Magdevska, M. Gradišek, E. Merljak, K. Cerović and Ž. Pušnik for their contribution in carrying out the initial experiments on which the rest of the work was based. Apart from the members of the 2016 iGEM team we would also like to thank B. Domevščik for helping with the experimental work. Lastly, we would also like to thank (Institute of Biotechnology, Swiss Federal Institute of Technology, ETH Zürich) for providing the plasmid p_PIR_ON-SEAP-pA.

## Author contributions

AP, TF, NF, JL, and RJ designed the experiments. AP, TF, NF and JL analyzed the data. NJ, TP and SR performed the experiments. AP and MB performed single cell analysis and protein trafficking. RJ conceived and supervised the study. AP, TF and RJ wrote the manuscript. All authors discussed and commented on manuscript.

## Competing interests

The authors declare no competing interests.

## Notes

### Competing Interest Statement

The authors have declared no competing interest.

