## Supplemental file for "Regulation of protein secretion through chemical regulation of endoplasmic reticulum retention signal cleavage"

**Supplementary Table 1:** Amino acid sequences of constructs used in this study. Unless stated otherwise, all constructs contained the CMV promoter and were inserted in the pcDNA3 backbone.

| **Potyviral proteases, ER-localized proteases and inducible split-proteases** | | |
|---|---|---|
| № | Name and scheme of construct | Amino acid or nucleotide sequence and description of parts |
| 1 | TEVp 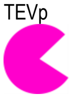 | M EQKLISEEDL GESLFKGPRDYNPISSTICHLTNESDGHTTSLYGIGFGPFIITNKHLFRRNNGTLLVQSLHGVFKVKNTTTLQQHLIDGRD<br>MIIIRMPKDFPPFPQKLKFREPQREERICLVTTNFQTKSMSSMVSDTSCTFPSSDGIFWKHWIQTKDGQCGSPLVSTRDGFIVGIHSASNFTNT<br>NNYFTSVPKNFMELLTNQEAQQWVSGWRLNADSVLWGGHKVFMSKPEEPFQPVKEATQLMSELVYSQYPYDVPDYA<br><br>Dark blue: Myc tag; Magenta: TEVp |
| 2 | PPVp 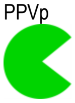 | M SKSLFRGLRDYNPIASSICQLNNSSGARQSEMFGLGFGGLIVTNQHLFKRNDGELTIRSHHGEFVVKDTKTLKLLPCKGRDIVIIRLPKDFP<br>PFPKRLQFRTPTTEDRVCLIGSNFQTKSISSTMSETSATYPVDNSHFWKHWISTKDGHCGLPIVSTRDGSILGLHSLANSTNTQNFYAAFPDNF<br>ETTYLSNQDNDNWIKQWRYNPDEVCWGSLQLKRDIPQSPFTICKLLTDLDGEFVYTQ<br><br>Greem: PPVp |
| 3 | SbMVp 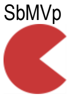 | M SKSVYKGLRDYSGISTLICQLTNSSDGHKETMFGVGYGSFIITNGHLFRRNNGMLTVKTWHGEFVIHNTTQLKIHFIQGRDVILIRMPKDFP<br>PFGKRNLFRQPKREERVCMVGTNFQEKSLRATVSESSMILPEGKGSFWIHWITTQDGFCGLPLVSVNDGHIVGIHGLTSNDSEKNFFVPLTDGF<br>EKEYLENADNLSWDKHWFWEPSKIAWGSLNLVEEQPKEEFKISKLVSDLFGNTVTVQ YPYDVPDYA<br><br>Dark brown: SbMVp; Dark blue: HA tag |
| 4 | SuMMVp 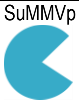 | M GVSLSRGVRDYNAISSMVCRVTNDSGSSSTTMYGIGYGCYIITNKHLFRENNGRLLITSHHGEYICKNSASLKLSLVPGRDMLLIRLPKDCP<br>PFPSKLKFREPTSEEKAVLVVTNFQEKHLSSMVSESSCVVQREDSPIWRHWISTKDGHCGAPIVSIRDGYIIGSHCGENPMTSNFFTSIPKDFQ<br>NLLNGKEANEWVSGWKYNIDAVCWGGLSVVNDAPSEPFITAKVVSALDTEGIKVQ YPYDVPDYA<br><br>Blue: SuMMVp; Dark blue: HA tag |
| 5 | erTEVp 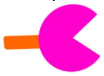 | M DMRVLAQLLGLLLLCFPGARCGESLFKGPRDYNPISSTICHLTQESDGHTTSLYGIGFGPFIITNKHLFRRNNGTLLVQSLHGVFKVKNTT<br>TLQQHLIDGRDMIIIRMPKDFPPFPQKLKFREPQREERICLVTTNFQTKSMSSMVSDTSSTFPSSDGIFWKHWIQTKDGQCGSPLVSTRDGFIV<br>GIHSASNFGNTNNYFTSVPKNFMELLTNQEAQQWVSGWRLNADSVLWGGHKVFMSKPEEPFQPVKEATQLMNEGGGLE KDEL<br><br>Gray: Signaling sequence; Magenta: TEVp; Orange: KDEL signal |
| 6 | erPPVp 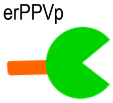 | M DMRVLAQLLGLLLLCFPGARCSKSLFRGLRDYNPIASSICQLNNSSGARQSEMFGLGFGGLIVTNQHLFKRNDGELTIRSHHGEFVVKDTK<br>TLKLLPCKGRDIVIIRLPKDFPPFPKRLQFRTPTTEDRVCLIGSNFQTKSISSTMSETSATYPVDNSHFWKHWISTKDGHCGLPIVSTRDGSIL<br>GLHSLANSTNTQNFYAAFPDNFETTYLSNQDNDNWIKQWRYNPDEVCWGSLQLKRDIPQSPFTICKLLTDLDGEFVYTQ KDEL<br><br>Gray: Signaling sequence; Green: PPVp; Orange: KDEL signal |
| 7 | erPPVp(N23Q, T173G) 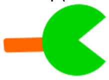 | M DMRVLAQLLGLLLLCFPGARCSKSLFRGLRDYNPIASSICQLNQSSGARQSEMFGLGFGGLIVTNQHLFKRNDGELTIRSHHGEFVVKDTKT<br>LKLLPCKGRDIVIIRLPKDFPPFPKRLQFRTPTTEDRVCLIGSNFQTKSISSTMSETSATYPVDNSHFWKHWISTKDGHCGLPIVSTRDGSILG<br>LHSLANSGNTQNFYAAFPDNFETTYLSNQDNDNWIKQWRYNPDEVCWGSLQLKRDIPQSPFTICKLLTDLDGEFVYTQ KDEL<br><br>Gray: Signaling sequence; Green: PPVp; Orange: KDEL signal |
| 8 | erSbMVp 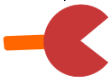 | M DMRVLAQLLGLLLLCFPGARCSKSVYKGLRDYSGISTLICQLTNSSDGHKETMFGVGYGSFIITNGHLFRRNNGMLTVKTWHGEFVIHNTTQ<br>LKIHFIQGRDVILIRMPKDFPPFGKRNLFRQPKREERVCMVGTNFQEKSLRATVSESSMILPEGKGSFWIHWITTQDGFCGLPLVSVNDGHIVG<br>IHGLTSNDSEKNFFVPLTDGFEKEYLENADNLSWDKHWFWEPSKIAWGSLNLVEEQPKEEFKISKLVSDLFGNTVTVQ KDEL<br><br>Gray: Signaling sequence; Dark brown: SbMVp; Orange: KDEL signal |
| 9 | ABI_cTEVp 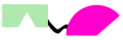 | M EQKLISEED LTRVPLYGFTSICGRRPEMEAAVSTIPRFLQSSSGSMLDGRFDPQSAAHFFGVYDGHGGSQVANYCRERMHLALA<br>EEIAKEKPMLCDGDTWLEKWKKALFNSFLRVDSEIESVAPETVGSTSVVAVVFPSHIFVANCGDSRAVLCRGKTALPLSVDHKPDREDEAARIE<br>AAGGKVIQWNGARVFGVLAMSRSIGDRYLKPSIIPDPEVTAVKRVKEDDCLILASDGVWDVMTDEEACEMARKRILLWHKKNAVAGDASLLADE<br>RRKEGKDPAAMSAAEYLSKLAIQRGSKDNISVVVVDLKGSGSKSMSSMVSDTSCTFPSSDGIFWKHWIQTKDGQCGSPLVSTRDGFIVGIHSAS<br>NFTNTNNYFTSVPKNFMELLTNQEAQQWVSGWRLNADSVLWGGHKVFMSKPEEPFQPVKEATQLMSELVYSQ<br><br>Dark blue: Myc tag; Black: ABI; Magenta: cTEVp |
| 10 | PYL1_nTEVp 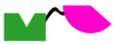 | M DTYRYI GGGAPTQDEFTQLSQSIAEFHTYQLGNGRCSSLLAQRIHAPPETVWSVVRRFDRPQIYKHFIKSCNVSEDFEMRVGCT<br>RDVNVISGLPANTSRERLDLLDDDRRVTGFSITGGEHRLRNYKSVTTVHRFEKEEEEERIWTVVLESYVVDVPEGNSEEDTRLFADTVIRLNLQ<br>KLASITEAMN GSGSS GESLFKGPRDYNPISSTICHLTNESDGHTTSLYGIGFGPFIITNKHLFRRNNGTLLVQSLHGV<br>FKVKNTTTLQQHLIDGRDMIIIRMPKDFPPFPQKLKFREPQREERICLVTTNFQT<br><br>Dark blue: AU1 tag; Black: PYL1; Magenta: nTEVp |
| 11 | FRB_nTEVp 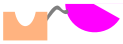 | M EQKLISEEDL ILWHEMWHEGLEEASRLYFGERNVKGMFEVLEPLHAMMERGPQTLKETSFNQAYGRDLMEAQEWCRKYMKSGNVKDLLQAW<br>DLYYHVFRRISK GSGS GESLFKGPRDYNPISSTICHLTNESDGHTTSLYGIGFGPFIITNKHLFRRNNGTLLVQSLHGVFKVKNTTTLQQHL<br>IDGRDMIIIRMPKDFPPFPQKLKFREPQREERICLVTTNFQT<br><br>Dark blue: Myc tag; Black: FRB; Magenta: nTEVp |
| 11 | FKBP_cTEVp 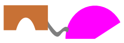 | M GVQVETISPGDGRTFPKRGQTCVVHYTGMLEDGKKFDSSRDRNKPFKFMLGKQEVIRGWEEGVAQMSVGQRAKLTISPDYAYGATGHPGIIP<br>PHATLVFDVELLKLE GSG KSMSSMVSDTSCTFPSSDGIFWKHWIQTKDGQCGSPLVSTRDGFIVGIHSASNFTNTNNYFTSVPKNFMELLTN<br>QEAQQWVSGWRLNADSVLWGGHKVFMSKPEEPFQPVKEATQLMSELVYSQ<br><br>Black: FKBP; Magenta: cTEVp; |
| 13 | FRB_NerTEVp 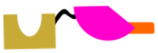 | M DMRVLAQLLGLLLLCFPGARC ILWHEMWHEGLEEASRLYFGERNVKGMFEVLEPLHAMMERGPQTLKETSFNQAYGRDLMEAQEWCRKYMK<br>SGNVKDLLQAWDLYYHVFRRISKGSGSGESLFKGPRDYNPISSTICHLTQESDGHTTSLYGIGFGPFIITNKHLFRRNNGTLLVQSLHGVFKVK<br>NTTTLQQHLIDGRDMIIIRMPKDFPPFPQKLKFREPQREERICLVTTNFQT KDEL<br><br>Gray: SS; Black: FRB; Magenta: nTEVp; Orange: KDEL signal |
| 14 | FKBP_CerTEVp | M DMRVLAQLLGLLLLCFPGARC<br>GVQVETISPGDGRTFPKRGQTCVVHYTGMLEDGKKFDSSRDRNKPFKFMLGKQEVIRGWEEGVAQMSVGQRAKLTISPDYAYGATGHPGIIPPH<br>ATLVFDVELLKLE GSG KSMSSMVSDTSSTFPSSDGIFWKHWIQTKDGQCGSPLVSTRDGFIVGIHSASNFGNTNNYFTSVPKNFMELLTNQE<br>AQQWVSGWRLNADSVLWGGHKVFMSKPEEPFQPVKEATQLMNEGGGLE KDEL |

| | | |
|---|---|---|
| | 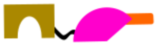 | Gray: SS;Black: FKBP; Magenta: cTEVp; Orange: KDEL signal |
| 15 | ABI-CerTEVp<br>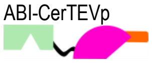 | M DMRVLAQLLGLLLLCFPGARC LTRVPLYGFTSICGRRPEMEAAVSTIPRFLQSSSGSMLDGRFDPQSAAHFFGVYDGHGGSQVANYCRERM<br>HLALA<br>EEIAKEKPMLCDGDTWLEKWKKALFNSFLRVDSEIESVAPETVGSTSVVAVVFPSHIFVANCGDSRAVLCRGKTALPLSVDHKPDREDEAARIE<br>AAGGKVIQWNGARVFGVLAMSRSIGDRYLKPSIIPDPEVTAVKRVKEDDCLILASDGVWDVMTDEEACEMARKRILLWHKKNAVAGDASLLADE<br>RRKEGKDPAAMSAAEYLSKLAIQRGSKDNISVVVVDLK<br>GSG KSMSSMVSDTSSTFPSSDGIFWKHWIQTKDGQCGSPLVSTRDGFIVGIHSASNFGNTNNYFTSVPKNFMELLTNQEAQQWVSGWRLNADS<br>VLWGGHKVFMSKPEEPFQPVKEATQLMNEGGGLE KDEL<br><br>Gray: SS; Black: ABI; Magenta: cTEVp Orange: KDEL signal |
| 16 | PYL1-NerTEVp<br>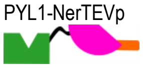 | M DMRVLAQLLGLLLLCFPGARC GGGAPTQDEFTQLSQSIAEFHTYQLGNGRCSSLLAQRIHAPPETVWSVVRRFDRPQIYKHFIKSCNVSED<br>FEMRVGCT<br>RDVNVISGLPANTSRERLDLLDDDRRVTGFSITGGEHRLRNYKSVTTVHRFEKEEEEERIWTVVLESYVVDVPEGNSEEDTRLFADTVIRLNLQ<br>KLASITEAMN GSGS<br>GESLFKGPRDYNPISSTICHLTQESDGHTTSLYGIGFGPFIITNKHLFRRNNGTLLVQSLHGVFKVKNTTTLQQHLIDGRDMIIIRMPKDFPPF<br>PQKLKFREPQREERICLVTTNFQT KDEL<br><br>Gray: SS; Black: PYL1; Magenta: nTEVp; Orange: KDEL signal |
| 17 | ABI-cPPVp<br>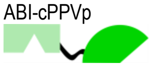 | M EQKLISEED LTRVPLYGFTSICGRRPEMEAAVSTIPRFLQSSSGSMLDGRFDPQSAAHFFGVYDGHGGSQVANYCRERMHLALA<br>EEIAKEKPMLCDGDTWLEKWKKALFNSFLRVDSEIESVAPETVGSTSVVAVVFPSHIFVANCGDSRAVLCRGKTALPLSVDHKPDREDEAARIE<br>AAGGKVIQWNGARVFGVLAMSRSIGDRYLKPSIIPDPEVTAVKRVKEDDCLILASDGVWDVMTDEEACEMARKRILLWHKKNAVAGDASLLADE<br>RRKEGKDPAAMSAAEYLSKLAIQRGSKDNISVVVVDLK GSGS<br>KSISSTMSETSATYPVDNSHFWKHWISTKDGHCGLPIVSTRDGSILGLHSLANSTNTQNFYAAFPDNFETTYLSNQDNDNWIKQWRYNPDEVCW<br>GSLQLKRDIPQSPFTICKLLTDLDGEFVYTQ<br><br>Dark blue: Myc tag; Black: ABI; Green: cPPVp |
| 18 | PYL-nPPVp<br>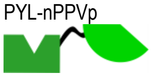 | M DTYRYI GGGAPTQDEFTQLSQSIAEFHTYQLGNGRCSSLLAQRIHAPPETVWSVVRRFDRPQIYKHFIKSCNVSEDFEMRVGCT<br>RDVNVISGLPANTSRERLDLLDDDRRVTGFSITGGEHRLRNYKSVTTVHRFEKEEEEERIWTVVLESYVVDVPEGNSEEDTRLFADTVIRLNLQ<br>KLASITEAMN GSGS<br>SKSLFRGLRDYNPIASSICQLNNSSGARQSEMFGLGFGGLIVTNQHLFKRNDGELTIRSHHGEFVVKDTKTLKLLPCKGRDIVIIRLPKDFP<br>PFPKRLQFRTPTTEDRVCLIGSNFQT<br><br>Dark blue: AU1 tag; Black: PYL1; Green: nPPVp |
| 19 | FKBP_PPVs_cTEVp<br>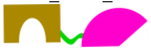 | M GVQVETISPGDGRTFPKRGQTCVVHYTGMLEDGKKFDSSRDRNKPFKFMLGKQEVIRGWEEGVAQMSVGQRAKLTISPDYAYGATGHPGIIP<br>PHATLVFDVELLKLE GSG NVVVHQA GSG<br>KSMSSMVSDTSCTFPSSDGIFWKHWIQTKDGQCGSPLVSTRDGFIVGIHSASNFTNTNNYFTSVPKNFMELLTNQEAQQWVSGWRLNADSVLWG<br>GHKVFMSKPEEPFQPVKEATQLMSELVYSQ<br><br>Black: FKBP; Green: PPVs; Magenta: cTEVp; |
| 20 | FRB_PPVs-nTEVp<br>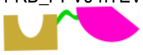 | M EQKLISEEDL ILWHEMWHEGLEEASRLYFGERNVKGMFEVLEPLHAMMERGPQTLKETSFNQAYGRDLMEAQEWCRKYMKSGNVKDLLQAW<br>DLYYHVFRRISK GSG NVVVHQA GSG<br>GESLFKGPRDYNPISSTICHLTNESDGHTTSLYGIGFGPFIITNKHLFRRNNGTLLVQSLHGVFKVKNTTTLQQHLIDGRDMIIIRMPKDFPPF<br>PQKLKFREPQREERICLVTTNFQT<br><br>Dark blue: Myc tag; Black: FRB; Green: PPVs; Magenta: nTEVp |

**membER and lumER secretion constructs and reporters**

| № | Name and scheme of construct | Amino acid or nucleotide sequence and description of parts |
|---|---|---|
| 21 | SEAP<br>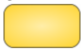 | M LLLLLLLGLRLQLSLGIIPVEEENPDFWNREAAEALGAAKKLQPAQTAAKNLIIFLGDGMGVSTVTAARILKGQKKDKLGPEIPL<br>AMDRFPYVALSKTYNVDKHVPDSGATATAYLCGVKGNFQTIGLSAAARFNQCNTTRGNEVISVMNRAKKAGKSVGVVTTTRVQHASPAGTYAHT<br>VNRNWYSDADVPASARQEGCQDIATQLISNMDIDVILGGGRKYMFRMGTPDPEYPDDYSQGGTRLDGKNLVQEWLAKRQGARYVWNRTELMQAS<br>LDPSVTHLMGLFEPGDMKYEIHRDSTLDPSLMEMTEAALRLLSRNPRGFFLFVEGGRIDHGHHESRAYRALTETIMFDDAIERAGQLTSEEDTL<br>SLVTADHSHVFSFGGYPLRGSSIFGLAPGKARDRKAYTVLLYGNGPGYVLKDGARPDVTESESGSPEYRQQSAVPLDEETHAGEDVAVFARGPQ<br>AHLVHGVQEQTFIAHVMAFAACLEPYTACDLAPPAGTTDAAHPGYSRVGAAGRFEQT DTYRYIE<br><br>Gold: SEAP: Dark blue: AU1 tag; |
| 22 | SEAP_TEVs_KDEL<br>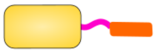 | M LLLLLLLGLRLQLSLGIIPVEEENPDFWNREAAEALGAAKKLQPAQTAAKNLIIFLGDGMGVSTVTAARILKGQKKDKLGPEIPL<br>AMDRFPYVALSKTYNVDKHVPDSGATATAYLCGVKGNFQTIGLSAAARFNQCNTTRGNEVISVMNRAKKAGKSVGVVTTTRVQHASPAGTYAHT<br>VNRNWYSDADVPASARQEGCQDIATQLISNMDIDVILGGGRKYMFRMGTPDPEYPDDYSQGGTRLDGKNLVQEWLAKRQGARYVWNRTELMQAS<br>LDPSVTHLMGLFEPGDMKYEIHRDSTLDPSLMEMTEAALRLLSRNPRGFFLFVEGGRIDHGHHESRAYRALTETIMFDDAIERAGQLTSEEDTL<br>SLVTADHSHVFSFGGYPLRGSSIFGLAPGKARDRKAYTVLLYGNGPGYVLKDGARPDVTESESGSPEYRQQSAVPLDEETHAGEDVAVFARGPQ<br>AHLVHGVQEQTFIAHVMAFAACLEPYTACDLAPPAGTTDAAHPGYSRVGAAGRFEQT DTYRYIE ENLYFQS KDEL<br><br>Gold: SEAP: Dark blue: AU1 tag; Black: TEV protease cleavage site; Orange: KDEL signal |
| 23 | SEAP_TM_3xTEVs_KKMP<br>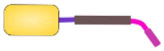 | M LLLLLLLGLRLQLSLGIIPVEEENPDFWNREAAEALGAAKKLQPAQTAAKNLIIFLGDGMGVSTVTAARILKGQKKDKLGPEIPL<br>AMDRFPYVALSKTYNVDKHVPDSGATATAYLCGVKGNFQTIGLSAAARFNQCNTTRGNEVISVMNRAKKAGKSVGVVTTTRVQHASPAGTYAHT<br>VNRNWYSDADVPASARQEGCQDIATQLISNMDIDVILGGGRKYMFRMGTPDPEYPDDYSQGGTRLDGKNLVQEWLAKRQGARYVWNRTELMQAS<br>LDPSVTHLMGLFEPGDMKYEIHRDSTLDPSLMEMTEAALRLLSRNPRGFFLFVEGGRIDHGHHESRAYRALTETIMFDDAIERAGQLTSEEDTL<br>SLVTADHSHVFSFGGYPLRGSSIFGLAPGKARDRKAYTVLLYGNGPGYVLKDGARPDVTESESGSPEYRQQSAVPLDEETHAGEDVAVFARGPQ<br>AHLVHGVQEQTFIAHVMAFAACLEPYTACDLAPPAGTTDAAHPGYSRVGAAGRFEQT<br>DTYRYI EARNRQKR GSGS ARNRQKR GSGS ARNRQKR SGS IIMIQTLLIILFIIVPIFLL SGSG ENLYFQS GSG ENLYFQS<br>GSG ENLYFQS GSG KKMP<br><br>Gold: SEAP: Dark blue: AU1 tag; FAS cleavage site; Purple: transmembrane domain; Cyan: TEV protease cleavage site; Orange: KKMP signal |
| 24 | SEAP_PPVs_KDEL<br>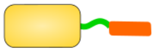 | M LLLLLLLGLRLQLSLGIIPVEEENPDFWNREAAEALGAAKKLQPAQTAAKNLIIFLGDGMGVSTVTAARILKGQKKDKLGPEIPL<br>AMDRFPYVALSKTYNVDKHVPDSGATATAYLCGVKGNFQTIGLSAAARFNQCNTTRGNEVISVMNRAKKAGKSVGVVTTTRVQHASPAGTYAHT<br>VNRNWYSDADVPASARQEGCQDIATQLISNMDIDVILGGGRKYMFRMGTPDPEYPDDYSQGGTRLDGKNLVQEWLAKRQGARYVWNRTELMQAS<br>LDPSVTHLMGLFEPGDMKYEIHRDSTLDPSLMEMTEAALRLLSRNPRGFFLFVEGGRIDHGHHESRAYRALTETIMFDDAIERAGQLTSEEDTL |

| | | |
|---|---|---|
| | | SLVTADHSHVFSFGGYPLRGSSIFGLAPGKARDRKAYTVLLYGNGPGYVLKDGARPDVTESESGSPEYRQQSAVPLDEETHAGEDVAVFARGPQ<br>AHLVHGVQEQTFIAHVMAFAACLEPYTACDLAPPAGTTDAAHPGYSRVGAAGRFEQT NVVVHQA KDEL<br>Gold: SEAP: Black: PPV protease cleavage site; Orange: KDEL signal |
| 25 | SEAP_SbMVs_KDEL<br>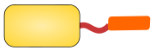 | M LLLLLLLGLRLQLSLGIIPVEEENPDFWNREAAEALGAAKKLQPAQTAAKNLIIFLGDGMGVSTVTAARILKGQKKDKLGPEIPL<br>AMDRFPYVALSKTYNVDKHVPDSGATATAYLCGVKGNFQTIGLSAAARFNQCNTTRGNEVISVMNRAKKAGKSVGVVTTTRVQHASPAGTYAHT<br>VNRNWYSDADVPASARQEGCQDIATQLISNMDIDVILGGGRKYMFRMGTPDPEYPDDYSQGGTRLDGKNLVQEWLAKRQGARYVWNRTELMQAS<br>LDPSVTHLMGLFEPGDMKYEIHRDSTLDPSLMEMTEAALRLLSRNPRGFFLFVEGGRIDHGHHESRAYRALTETIMFDDAIERAGQLTSEEDTL<br>SLVTADHSHVFSFGGYPLRGSSIFGLAPGKARDRKAYTVLLYGNGPGYVLKDGARPDVTESESGSPEYRQQSAVPLDEETHAGEDVAVFARGPQ<br>AHLVHGVQEQTFIAHVMAFAACLEPYTACDLAPPAGTTDAAHPGYSRVGAAGRFEQT ESVSLQS KDEL<br>Gold: SEAP: Black: SbMV protease cleavage site; Orange: KDEL signal |
| 26 | SEAP_AU1_3xFURS_TM_<br>3xTEVs_KKYL<br>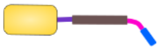 | M LLLLLLLGLRLQLSLGIIPVEEENPDFWNREAAEALGAAKKLQPAQTAAKNLIIFLGDGMGVSTVTAARILKGQKKDKLGPEIPL<br>AMDRFPYVALSKTYNVDKHVPDSGATATAYLCGVKGNFQTIGLSAAARFNQCNTTRGNEVISVMNRAKKAGKSVGVVTTTRVQHASPAGTYAHT<br>VNRNWYSDADVPASARQEGCQDIATQLISNMDIDVILGGGRKYMFRMGTPDPEYPDDYSQGGTRLDGKNLVQEWLAKRQGARYVWNRTELMQAS<br>LDPSVTHLMGLFEPGDMKYEIHRDSTLDPSLMEMTEAALRLLSRNPRGFFLFVEGGRIDHGHHESRAYRALTETIMFDDAIERAGQLTSEEDTL<br>SLVTADHSHVFSFGGYPLRGSSIFGLAPGKARDRKAYTVLLYGNGPGYVLKDGARPDVTESESGSPEYRQQSAVPLDEETHAGEDVAVFARGPQ<br>AHLVHGVQEQTFIAHVMAFAACLEPYTACDLAPPAGTTDAAHPGYSRVGAAGRFEQT<br>DTYRYIE ARNRQKR GSGS ARNRQKR GSGS ARNRQKR SGS IIMIQTLLIILFIIVPIFLL SGSG ENLYFQS GSG ENLYFQS<br>GSG ENLYFQS GSG KKYL<br><br>Gold: SEAP: Dark blue: AU1 tag; FAS cleavage site; Purple: transmembrane domain; Cyan: TEV protease cleavage site; Orange: KKYL signal |
| 27 | SEAP_AU1_3xFURS_TM_<br>3xPPVs_KKYL<br>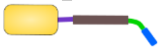 | M LLLLLLLGLRLQLSLGIIPVEEENPDFWNREAAEALGAAKKLQPAQTAAKNLIIFLGDGMGVSTVTAARILKGQKKDKLGPEIPL<br>AMDRFPYVALSKTYNVDKHVPDSGATATAYLCGVKGNFQTIGLSAAARFNQCNTTRGNEVISVMNRAKKAGKSVGVVTTTRVQHASPAGTYAHT<br>VNRNWYSDADVPASARQEGCQDIATQLISNMDIDVILGGGRKYMFRMGTPDPEYPDDYSQGGTRLDGKNLVQEWLAKRQGARYVWNRTELMQAS<br>LDPSVTHLMGLFEPGDMKYEIHRDSTLDPSLMEMTEAALRLLSRNPRGFFLFVEGGRIDHGHHESRAYRALTETIMFDDAIERAGQLTSEEDTL<br>SLVTADHSHVFSFGGYPLRGSSIFGLAPGKARDRKAYTVLLYGNGPGYVLKDGARPDVTESESGSPEYRQQSAVPLDEETHAGEDVAVFARGPQ<br>AHLVHGVQEQTFIAHVMAFAACLEPYTACDLAPPAGTTDAAHPGYSRVGAAGRFEQT<br>DTYRYIE ARNRQKR GSGS ARNRQKR GSGS ARNRQKR SGS IIMIQTLLIILFIIVPIFLL SGSG NVVVHQA GSG NVVVHQA<br>GSG NVVVHQA GSG KKYL<br><br>Gold: SEAP: Dark blue: AU1 tag; FAS cleavage site; Purple: transmembrane domain; Cyan: PPV protease cleavage site; Orange: KKYL signal |
| 28 | SEAP_AU1_3xFURS_TM_<br>3xSbMVs_KKYL<br>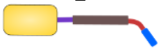 | M LLLLLLLGLRLQLSLGIIPVEEENPDFWNREAAEALGAAKKLQPAQTAAKNLIIFLGDGMGVSTVTAARILKGQKKDKLGPEIPL<br>AMDRFPYVALSKTYNVDKHVPDSGATATAYLCGVKGNFQTIGLSAAARFNQCNTTRGNEVISVMNRAKKAGKSVGVVTTTRVQHASPAGTYAHT<br>VNRNWYSDADVPASARQEGCQDIATQLISNMDIDVILGGGRKYMFRMGTPDPEYPDDYSQGGTRLDGKNLVQEWLAKRQGARYVWNRTELMQAS<br>LDPSVTHLMGLFEPGDMKYEIHRDSTLDPSLMEMTEAALRLLSRNPRGFFLFVEGGRIDHGHHESRAYRALTETIMFDDAIERAGQLTSEEDTL<br>SLVTADHSHVFSFGGYPLRGSSIFGLAPGKARDRKAYTVLLYGNGPGYVLKDGARPDVTESESGSPEYRQQSAVPLDEETHAGEDVAVFARGPQ<br>AHLVHGVQEQTFIAHVMAFAACLEPYTACDLAPPAGTTDAAHPGYSRVGAAGRFEQT<br>DTYRYIE ARNRQKR GSGS ARNRQKR GSGS ARNRQKR SGS IIMIQTLLIILFIIVPIFLL SGSG ESVSLQS GSG ESVSLQS<br>GSG ESVSLQS GSG KKYL<br><br>Gold: SEAP: Dark blue: AU1 tag; FAS cleavage site; Purple: transmembrane domain; Cyan: SbMV protease cleavage site; Orange: KKYL signal |
| 29 | SEAP_AU1_3xFURS_TM_<br>3xSuMMVs_KKYL<br>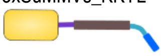 | M LLLLLLLGLRLQLSLGIIPVEEENPDFWNREAAEALGAAKKLQPAQTAAKNLIIFLGDGMGVSTVTAARILKGQKKDKLGPEIPL<br>AMDRFPYVALSKTYNVDKHVPDSGATATAYLCGVKGNFQTIGLSAAARFNQCNTTRGNEVISVMNRAKKAGKSVGVVTTTRVQHASPAGTYAHT<br>VNRNWYSDADVPASARQEGCQDIATQLISNMDIDVILGGGRKYMFRMGTPDPEYPDDYSQGGTRLDGKNLVQEWLAKRQGARYVWNRTELMQAS<br>LDPSVTHLMGLFEPGDMKYEIHRDSTLDPSLMEMTEAALRLLSRNPRGFFLFVEGGRIDHGHHESRAYRALTETIMFDDAIERAGQLTSEEDTL<br>SLVTADHSHVFSFGGYPLRGSSIFGLAPGKARDRKAYTVLLYGNGPGYVLKDGARPDVTESESGSPEYRQQSAVPLDEETHAGEDVAVFARGPQ<br>AHLVHGVQEQTFIAHVMAFAACLEPYTACDLAPPAGTTDAAHPGYSRVGAAGRFEQT<br>DTYRYIE ARNRQKR GSGS ARNRQKR GSGS ARNRQKR SGS IIMIQTLLIILFIIVPIFLL SGSG EEIHLQS GSG EEIHLQS<br>GSG EEIHLQS GSG KKYL<br><br>Gold: SEAP: Dark blue: AU1 tag; FAS cleavage site; Purple: transmembrane domain; Cyan: SuMMV protease cleavage site; Orange: KKYL signal |
| 30 | Gluc-TEVs-KDEL<br>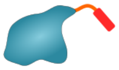 | M ALWMRLLPLLALLALWGPDPAAA KPTENNEDFNIVAVASNFATTDLDADRGKLPGKKLPLEVLKEMEANARKAGCTRGCLICLS<br>HIKCTPKMKKFIPGRCHTYEGDKESAQGGIGEAIVDIPEIPGFKDLEPMEQFIAQVDLCVDCTTGCLKGLANVQCSDLLKKWLPQRCATFASKI<br>QGQVDKIKGAGGD ENLYFQS KDEL<br><br>Gray: Signaling sequence; Blue: Gaussia luciferase; Black: TEV protease cleavage site; Orange: KDEL signal |
| 31 | fLuc<br> | M EQKLISEEDL EDAKNIKKGPAPFYPLEDGTAGEQLHKAMKRYALVPGTIAFTDAHIEVDITYAEYFEMSVRLAEAMKRYGLNTN<br>HRIVVCSENSLQFFMPVLGALFIGVAVAPANDIYNERELLNSMGISQPTVVFVSKKGLQKILNVQKKLPIIQKIIIMDSKTDYQGFQSMYTFVT<br>SHLPPGFNEYDFVPESFDRDKTIALIMNSSGSTGLPKGVALPHRTACVRFSHARDPIFGNQIIPDTAILSVVPFHHGFGMFTTLGYLICGFRVV<br>LMYRFEEELFLRSLQDYKIQSALLVPTLFSFFAKSTLIDKYDLSNLHEIASGGAPLSKEVGEAVAKRFHLPGIRQGYGLTETTSAILITPEGDD<br>KPGAVGKVVPFFEAKVVDLDTGKTLGVNQRGELCVRGPMIMSGYVNNPEATNALIDKDGWLHSGDIAYWDEDEHFFIVDRLKSLIKYKGYQVAP<br>AELESILLQHPNIFDAGVAGLPDDDAGELPAAVVVLEHGKSTMTEKEIVDYVASQVTTAKKLRGGVVFVDEVPKGLTGKLDARKIREILIKAKK<br>GGKIAVNSGSG YPYDVPDYA<br><br>Dark blue: Myc-tag; Black: fLuc; ; Blue: HA-tag |
| 32 | rLuc<br>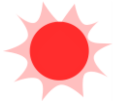 | M ASKVYDPEQRKRMITGPQWWARCKQMNVLDSFINYYDSEKHAENAVIFLHGNAASSYLWRHVVPHIEPVARCIIPDLIGMGKSGKSGNGSY<br>RLLDHYKYLTAWFELLNLPKKIIFVGHDWGACLAFHYSYEHQDKIKAIVHAESVVDVIESWDEWPDIEEDIALIKSEEGEKMVLENNFFVETML<br>PSKIMRKLEPEEFAAYLEPFKEKGEVRRPTLSWPREIPLVKGGKPDVVQIVRNYNAYLRASDDLPKMFIESDPGFFSNAIVEGAKKFPNTEFVK<br>VKGLHFSQEDAPDEMGKYIKSFVERVLKNEQ<br><br>Red: *Renilla* luciferase |

**Constructs for inducible transcription**

| № | Name and scheme of construct | Amino acid or nucleotide sequence and description of parts |
|---|---|---|
| 33 | PYL1_NLS_VPR<br>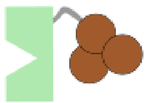 | M HHHHHH GGGAPTQDEFTQLSQSIAEFHTYQLGNGRCSSLLAQRIHAPPETVWSVVRRFDRPQIYKHFIKSCNVSEDFEMRVGCT<br>RDVNVISGLPANTSRERLDLLDDDRRVTGFSITGGEHRLRNYKSVTTVHRFEKEEEEERIWTVVLESYVVDVPEGNSEEDTRLFADTVIRLNLQ<br>KLASITEAMN DPKKKRKV EASGSGRADALDDFDLDMLGSDALDDFDLDMLGSDALDDFDLDMLGSDALDDFDLDMLIN<br>TSGGSGGGSGGSSQYLPDTDDRHRIEEKRKRTYETFKSIMKKSPFSGPTDPRPPPRRIAVPSRSSASVPKPAPQPYPFTSSLSTINYDEFPTMV<br>FPSGQISQASALAPAPPQVLPQAPAPAPAPAMVSALAQAPAPVPVLAPGPPQAVAPPAPKPTQAGEGTLSEALLQLQFDDEDLGALLGNSTDPA<br>VFTDLASVDNSEFQQLLNQGIPVAPHTTEPMLMEYPEAITRLVTGAQRPPDPAPAPLGAPGLPNGLLSGDEDFSSIADMDFSALLGSGSGSRDS |

| | | |
|---|---|---|
| | | REGMFLPKPEAGSAISDVFEGREVCQPKRIRPFHPPGSPWANRPLPASLAPTPTGPVHEPVGSLTPAPVPQPLDPAPAVTPEASHLLEDPDEET<br>SQAVKALREMADTVIPQKEEAAICGQMDLSHPPPRGHLDELTTTLESMTEDLNLDSPLTPELNEILDTFLNDECLLHAMHISTGLSIFDTSLF<br><br>Dark blue: His tag; Black: PYL1; Red: nuclear localization signal; Orange: transcription activation domain VPR |
| 34 | FRB_NLS_VPR<br>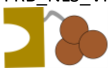 | M EQKLISEEDL ILWHEMWHEGLEEASRLYFGERNVKGMFEVLEPLHAMMERGPQTLKETSFNQAYGRDLMEAQEWCRKYMKSGNVKDLLQAW<br>DLYYHVFRRISK GSGS DPKKKRKV EASGSGRADALDDFDLDMLGSDALDDFDLDMLGSDALDDFDLDMLGSDALDDFDLDMLIN<br>TSGGSGGGSGGSSQYLPDTDDRHRIEEKRKRTYETFKSIMKKSPFSGPTDPRPPPRRIAVPSRSSASVPKPAPQPYPFTSSLSTINYDEFPTMV<br>FPSGQISQASALAPAPPQVLPQAPAPAPAPAMVSALAQAPAPVPVLAPGPPQAVAPPAPKPTQAGEGTLSEALLQLQFDDEDLGALLGNSTDPA<br>VFTDLASVDNSEFQQLLNQGIPVAPHTTEPMLMEYPEAITRLVTGAQRPPDPAPAPLGAPGLPNGLLSGDEDFSSIADMDFSALLGSGSGSRDS<br>REGMFLPKPEAGSAISDVFEGREVCQPKRIRPFHPPGSPWANRPLPASLAPTPTGPVHEPVGSLTPAPVPQPLDPAPAVTPEASHLLEDPDEET<br>SQAVKALREMADTVIPQKEEAAICGQMDLSHPPPRGHLDELTTTLESMTEDLNLDSPLTPELNEILDTFLNDECLLHAMHISTGLSIFDTSLF<br><br>Dark blue: Myc tag; Black: FRB; Red: nuclear localization signal; Orange: transcription activation domain VPR |
| 35 | dCas9_NLS_ABI<br>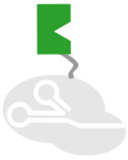 | M HHHHHH DKKYSIGLAIGTNSVGWAVITDEYKVPSKKFKVLGNTDRHSIKKNLIGALLFDSGETAEATRLKRTARRRYTRRKNRI<br>CYLQEIFSNEMAKVDDSFFHRLEESFLVEEDKKHERHPIFGNIVDEVAYHEKYPTIYHLRKKLVDSTDKADLRLIYLALAHMIKFRGHFLIEGD<br>LNPDNSDVDKLFIQLVQTYNQLFEENPINASGVDAKAILSARLSKSRRLENLIAQLPGEKKNGLFGNLIALSLGLTPNFKSNFDLAEDAKLQLS<br>KDTYDDDLDNLLAQIGDQYADLFLAAKNLSDAILLSDILRVNTEITKAPLSASMIKRYDEHHQDLTLLKALVRQQLPEKYKEIFFDQSKNGYAG<br>YIDGGASQEEFYKFIKPILEKMDGTEELLVKLNREDLLRKQRTFDNGSIPHQIHLGELHAILRRQEDFYPFLKDNREKIEKILTFRIPYYVGPL<br>ARGNSRFAWMTRKSEETITPWNFEEVVDKGASAQSFIERMTNFDKNLPNEKVLPKHSLLYEYFTVYNELTKVKYVTEGMRKPAFLSGEQKKAIV<br>DLLFKTNRKVTVKQLKEDYFKKIECFDSVEISGVEDRFNASLGTYHDLLKIIKDKDFLDNEENEDILEDIVLTLTLFEDREMIEERLKTYAHLF<br>DDKVMKQLKRRRYTGWGRLSRKLINGIRDKQSGKTILDFLKSDGFANRNFMQLIHDDSLTFKEDIQKAQVSGQGDSLHEHIANLAGSPAIKKGI<br>LQTVKVVDELVKVMGRHKPENIVIEMARENQTTQKGQKNSRERMKRIEEGIKELGSQILKEHPVENTQLQNEKLYLYYLQNGRDMYVDQELDIN<br>RLSDYDVDAIVPQSFLKDDSIDNKVLTRSDKNRGKSDNVPSEEVVKKMKNYWRQLLNAKLITQRKFDNLTKAERGGLSELDKAGFIKRQLVETR<br>QITKHVAQILDSRMNTKYDENDKLIREVKVITLKSKLVSDFRKDFQFYKVREINNYHHAHDAYLNAVVGTALIKKYPKLESEFVYGDYKVYDVR<br>KMIAKSEQEIGKATAKYFFYSNIMNFFKTEITLANGEIRKRPLIETNGETGEIVWDKGRDFATVRKVLSMPQVNIVKKTEVQTGGFSKESILPK<br>RNSDKLIARKKDWDPKKYGGFDSPTVAYSVLVVAKVEKGKSKKLKSVKELLGITIMERSSFEKNPIDFLEAKGYKEVKKDLIIKLPKYSLFELE<br>NGRKRMLASAGELQKGNELALPSKYVNFLYLASHYEKLKGSPEDNEQKQLFVEQHKHYLDEIIEQISEFSKRVILADANLDKVLSAYNKHRDKP<br>IREQAENIIHLFTLTNLGAPAAFKYFDTTIDRKRYTSTKEVLDATLIHQSITGLYETRIDLSQLGGD DPKKKRKV TRVPLY<br>GFTSICGRRPEMEAAVSTIPRFLQSSSGSMLDGRFDPQSAAHFFGVYDGHGGSQVANYCRERMHLALAEEIAKEKPMLCDGDTWLEKWKKALFN<br>SFLRVDSEIESVAPETVGSTSVVAVVFPSHIFVANCGDSRAVLCRGKTALPLSVDHKPDREDEAARIEAAGGKVIQWNGARVFGVLAMSRSIGD<br>RYLKPSIIPDPEVTAVKRVKEDDCLILASDGVWDVMTDEEACEMARKRILLWHKKNAVAGDASLLADERRKEGKDPAAMSAAEYLSKLAIQRGS<br>KDNISVVVVDLK<br><br>Dark blue: His tag; Orange: dCas9; Red: nuclear localization signal; Black: ABI |
| 36 | dCas9_NLS_FKBP<br>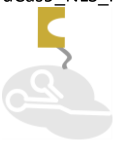 | M HHHHHH DKKYSIGLAIGTNSVGWAVITDEYKVPSKKFKVLGNTDRHSIKKNLIGALLFDSGETAEATRLKRTARRRYTRRKNRI<br>CYLQEIFSNEMAKVDDSFFHRLEESFLVEEDKKHERHPIFGNIVDEVAYHEKYPTIYHLRKKLVDSTDKADLRLIYLALAHMIKFRGHFLIEGD<br>LNPDNSDVDKLFIQLVQTYNQLFEENPINASGVDAKAILSARLSKSRRLENLIAQLPGEKKNGLFGNLIALSLGLTPNFKSNFDLAEDAKLQLS<br>KDTYDDDLDNLLAQIGDQYADLFLAAKNLSDAILLSDILRVNTEITKAPLSASMIKRYDEHHQDLTLLKALVRQQLPEKYKEIFFDQSKNGYAG<br>YIDGGASQEEFYKFIKPILEKMDGTEELLVKLNREDLLRKQRTFDNGSIPHQIHLGELHAILRRQEDFYPFLKDNREKIEKILTFRIPYYVGPL<br>ARGNSRFAWMTRKSEETITPWNFEEVVDKGASAQSFIERMTNFDKNLPNEKVLPKHSLLYEYFTVYNELTKVKYVTEGMRKPAFLSGEQKKAIV<br>DLLFKTNRKVTVKQLKEDYFKKIECFDSVEISGVEDRFNASLGTYHDLLKIIKDKDFLDNEENEDILEDIVLTLTLFEDREMIEERLKTYAHLF<br>DDKVMKQLKRRRYTGWGRLSRKLINGIRDKQSGKTILDFLKSDGFANRNFMQLIHDDSLTFKEDIQKAQVSGQGDSLHEHIANLAGSPAIKKGI<br>LQTVKVVDELVKVMGRHKPENIVIEMARENQTTQKGQKNSRERMKRIEEGIKELGSQILKEHPVENTQLQNEKLYLYYLQNGRDMYVDQELDIN<br>RLSDYDVDAIVPQSFLKDDSIDNKVLTRSDKNRGKSDNVPSEEVVKKMKNYWRQLLNAKLITQRKFDNLTKAERGGLSELDKAGFIKRQLVETR<br>QITKHVAQILDSRMNTKYDENDKLIREVKVITLKSKLVSDFRKDFQFYKVREINNYHHAHDAYLNAVVGTALIKKYPKLESEFVYGDYKVYDVR<br>KMIAKSEQEIGKATAKYFFYSNIMNFFKTEITLANGEIRKRPLIETNGETGEIVWDKGRDFATVRKVLSMPQVNIVKKTEVQTGGFSKESILPK<br>RNSDKLIARKKDWDPKKYGGFDSPTVAYSVLVVAKVEKGKSKKLKSVKELLGITIMERSSFEKNPIDFLEAKGYKEVKKDLIIKLPKYSLFELE<br>NGRKRMLASAGELQKGNELALPSKYVNFLYLASHYEKLKGSPEDNEQKQLFVEQHKHYLDEIIEQISEFSKRVILADANLDKVLSAYNKHRDKP<br>IREQAENIIHLFTLTNLGAPAAFKYFDTTIDRKRYTSTKEVLDATLIHQSITGLYETRIDLSQLGGD DPKKKRKV GVQVET<br>ISPGDGRTFPKRGQTCVVHYTGMLEDGKKFDSSRDRNKPFKFMLGKQEVIRGWEEGVAQMSVGQRAKLTISPDYAYGATGHPGIIPPHATLVFD<br>VELLKLE<br><br>Dark blue: His tag; Orange: dCas9; Red: nuclear localization signal; Black: FKBP |
| 37 | sgRNA [ab nt]<br>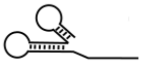 | AGATGTGGAAACGGAAGAAGC GTTTTAGAGCTAGAAATAGCAAGTTAAAATAAGGCTAGTCCGTTATCAACTTGAAAAAGTGGCAC<br>CGAGTCGGTGCTTTTTT<br><br>Orange: sgRNA targeting the nontemplate strand of sequence [ab]; Black: sgRNA scaffold |
| 38 | sgRNA [b]<br>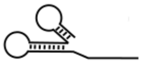 | TCTTCCGTTTCCACATCTTCA GTTTTAGAGCTAGAAATAGCAAGTTAAAATAAGGCTAGTCCGTTATCAACTTGAAAAAGTGGCAC<br>CGAGTCGGTGCTTTTTT<br><br>Orange: sgRNA targeting sequence [b]; Black: sgRNA scaffold |

### Constructs for microscopy

| № | Name and scheme of construct | Amino acid or nucleotide sequence and description of parts |
|---|---|---|
| 39 | TagRFP_FURS_TM_3xTEVs_KKYL<br>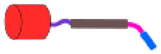 | M DMRVLAQLLGLLLLCFPGARC VSKGEELIKENMHMKLYMEGTVNNHHFKCTSEGEGKPYEGTQTMRIKVVEGGPLPFAFDILA<br>TSFMYGSRTFINHTQGIPDFFKQSFPEGFTWERVTTYEDGGVLTATQDTSLQDGCLIYNVKIRGVNFPSNGPVMQKKTLGWEANTEMLYPADGG<br>LEGRSDMALKLVGGGHLICNFKTTYRSKKPAKNLKMPGVYYVDHRLERIKEADKETYVEQHEVAVARYCDLPSKLGHKLN SARNRQKR<br>GSGSGG IIMIQTLLIILFIIVPIFLL SGSG ENLYFQS GSG ENLYFQS GSG ENLYFQS GSG KKYL<br><br>Gray: signaling sequence; Red: TagRFP; Cyan: furin protease cleavage site; Green: transmembrane domain; Magenta: TEV cleavage site; Orange: KKYL signal |
| 40 | TagRFP_FURS_TM_3xTEVs | M DMRVLAQLLGLLLLCFPGARC VSKGEELIKENMHMKLYMEGTVNNHHFKCTSEGEGKPYEGTQTMRIKVVEGGPLPFAFDILAT<br>SFMYGSRTFINHTQGIPDFFKQSFPEGFTWERVTTYEDGGVLTATQDTSLQDGCLIYNVKIRGVNFPSNGPVMQKKTLGWEANTEMLYPADGGL<br>EGRSDMALKLVGGGHLICNFKTTYRSKKPAKNLKMPGVYYVDHRLERIKEADKETYVEQHEVAVARYCDLPSKLGHKLN SARNRQKR<br>GSGSGG IIMIQTLLIILFIIVPIFLL SGSG ENLYFQS GSG ENLYFQS GSG ENLYFQS GSG |

| | | |
|---|---|---|
| | 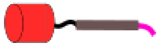 | Gray: signaling sequence; Red: TagRFP; Cyan: furin protease cleavage site; Green: transmembrane domain; Magenta: TEV cleavage site |
| 41 | TagRFP_AU1_3xFURS_TM_3xTEVs_KKMP<br>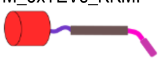 | M DMRVLAQLLGLLLLCFPGARC VSKGEELIKENMHMKLYMEGTVNNHHFKCTSEGEGKPYEGTQTMRIKVVEGGPLPFAFDILAT<br>SFMYGSRTFINHTQGIPDFFKQSFPEGFTWERVTTYEDGGVLTATQDTSLQDGCLIYNVKIRGVNFPSNGPVMQKKTLGWEANTEMLYPADGGL<br>EGRSDMALKLVGGGHLICNFKTTYRSKKPAKNLKMPGVYYVDHRLERIKEADKETYVEQHEVAVARYCDLPSKLGHKLN DTYRYIE<br>SARNRQKR GSGSGG IIMIQTLLIILFIIVPIFLL SGSG ENLYFQS GSG ENLYFQS GSG ENLYFQS GSG KKMP<br><br>Gray: signaling sequence; Red: TagRFP; Cyan: furin protease cleavage site; Green: transmembrane domain; Magenta: TEV cleavage site; Orange: KKMP signal |
| 42 | Ss_TagRFP_TEVS_KDEL<br>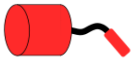 | M DMRVLAQLLGLLLLCFPGARC VSKGEELIKENMHMKLYMEGTVNNHHFKCTSEGEGKPYEGTQTMRIKVVEGGPLPFAFDILATSFMYGSR<br>TFINHTQGIPDFFKQSFPEGFTWERVTTYEDGGVLTATQDTSLQDGCLIYNVKIRGVNFPSNGPVMQKKTLGWEANTEMLYPADGGLEGRSDMA<br>LKLVGGGHLICNFKTTYRSKKPAKNLKMPGVYYVDHRLERIKEADKETYVEQHEVAVARYCDLPSKLGHKLNDTYRYIE ENLYFQS KDEL<br><br>Black: signal sequence; Red:TagRFP; Purple:AU1 tag; Magenta: TEV protease cleavage site; Orange: KDEL signal |
| 43 | TagRFP_AU1_TM_3xTEVs_KKYL<br>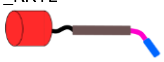 | M DMRVLAQLLGLLLLCFPGARC VSKGEELIKENMHMKLYMEGTVNNHHFKCTSEGEGKPYEGTQTMRIKVVEGGPLPFAFDILA<br>TSFMYGSRTFINHTQGIPDFFKQSFPEGFTWERVTTYEDGGVLTATQDTSLQDGCLIYNVKIRGVNFPSNGPVMQKKTLGWEANTEMLYPADGG<br>LEGRSDMALKLVGGGHLICNFKTTYRSKKPAKNLKMPGVYYVDHRLERIKEADKETYVEQHEVAVARYCDLPSKLGHKLN<br>IIMIQTLLIILFIIVPIFLL SGSG ENLYFQS GSG ENLYFQS GSG ENLYFQS GSG KKYL<br><br>Gray: signaling sequence; Red: TagRFP; Dark blue: AU1 tag; Green: transmembrane domain; Magenta: TEV cleavage site; Orange: KKYL signal |
| 44 | GFP1-10_TEVs_KDEL<br>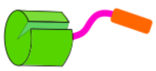 | M DMRVLAQLLGLLLLCFPGARC SKGEELFTGVVPILVELDGDVNGHKFSVRGEGEGDATIGKLTLKFICTTGKLPVPWPTLVTTL<br>TYGVQCFSRYPDHMKRHDFFKSAMPEGYVQERTISFKDDGKYKTRAVVKFEGDTLVNRIELKGTDFKEDGNILGHKLEYNFNSHNVYITADKQK<br>NGIKANFTVRHNVEDGSVQLADHYQQNTPIGDGPVLLPDNHYLSTQTVLSKDPNEK DTYRYI ENLYFQS KDEL<br><br>Gray: signaling sequence; Green: GFP1-10; Dark blue: AU1; Black: TEV protease cleavage site; Orange: KDEL signal |
| 45 | GFP1-10_3xFURs_TM_3xTEVs_KKYL<br>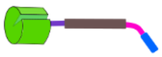 | M DMRVLAQLLGLLLLCFPGARC SKGEELFTGVVPILVELDGDVNGHKFSVRGEGEGDATIGKLTLKFICTTGKLPVPWPTLVTTL<br>TYGVQCFSRYPDHMKRHDFFKSAMPEGYVQERTISFKDDGKYKTRAVVKFEGDTLVNRIELKGTDFKEDGNILGHKLEYNFNSHNVYITADKQK<br>NGIKANFTVRHNVEDGSVQLADHYQQNTPIGDGPVLLPDNHYLSTQTVLSKDPNEK ARNRQKR GSGS ARNRQKR GSGS ARNRQKR SGS<br>IIMIQTLLIILFIIVPIFLL SGSG ENLYFQS GSG ENLYFQS GSG ENLYFQS GSG KKYL<br><br>Gray: signaling sequence; Light green: GFP1-10; Dark green: FAS cleavage site; Purple: transmembrane domain; Cyan: TEV protease cleavage site; Orange: KKYL signal |
| 46 | SS_GFP1-10_GS10_FASR<br>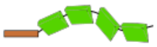 | M DMRVLAQLLGLLLLCFPGARC SKGEELFTGVVPILVELDGDVNGHKFSVRGEGEGDATIGKLTLKFICTTGKLPVPWPTLVTTL<br>TYGVQCFSRYPDHMKRHDFFKSAMPEGYVQERTISFKDDGKYKTRAVVKFEGDTLVNRIELKGTDFKEDGNILGHKLEYNFNSHNVYITADKQK<br>NGIKANFTVRHNVEDGSVQLADHYQQNTPIGDGPVLLPDNHYLSTQTVLSKDPNEK GGSGGGSGGS QVTDINSKGLEL<br>RKTVTTVETQNLEGLHHDGQFCHKPCPPGERKARDCTVNGDEPDCVPCQEGKEYTDKAHFSSKCRRCRLCDEGHGLEVEINCTRTQNTKCRCKP<br>NFFCNSTVCEHCDPCTKCEHGIIKECTLTSNTKCKEEGSRSNLGWLCLLLLPIPLIVWVKRKEVQKTCRKHRKENQGSHESPTLNPETVAINL<br><br>Gray: signaling sequence; Light green: GFP1-10; Orange: Fas receptor |
| 47 | iRFP<br>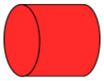 | M ARKVDLTSCDREPIHIPGSIQPCGCLLACDAQAVRITRITENAGAFFGRETPRVGELLADYFGETEAHALRNALAQSSDPKRPAL<br>IFGWRDGLTGRTFDISLHRHDGTSIIEFEPAAAEQADNPLRLTRQIIARTKELKSLEEMAARVPRYLQAMLGYHRVMLYRFADDGSGMVIGEAK<br>RSDLESFLGQHFPASLVPQQARLLYLKNAIRVVSDSRGISSRIVPEHDASGAALDLSFAHLRSISPCHLEFLRNMGVSASMSLSIIIDGTLWGL<br>IICHHYEPRAVPMAQRVAAEMFADFLSLHFTAAHHQR<br><br>Magenta: iRFP |
| 48 | BFP<br>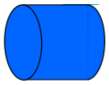 | M SELIKENMHMKLYMEGTVDNHHFKCTSEGEGKPYEGTQTMRIKVVEGGPLPFAFDILATSFLYGSKTFINHTQGIPDFFKQSFPE<br>GFTWERVTTYEDGGVLTATQDTSLQDGCLIYNVKIRGVNFTSNGPVMQKKTLGWEAFTETLYPADGGLEGRNDMALKLVGGSHLIANIKTTYRS<br>KKPAKNLKMPGVYYVDYRLERIKEANNETYVEQHEVAVARYCDLPSKLGHKLN<br><br>Black: BFP |
| 49 | Fas-EnvZ(C)-mCit<br>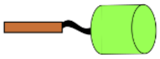 | M LGIWTLLPLVLTSVARLSSKSVNAYPYDVPDYAQVTDINSKGLELRKTVTTVETQNLEGLHHDGQFCHKPCPPGERKARDCTVN<br>GDEPDCVPCQEGKEYTDKAHFSSKCRRCRLCDEGHGLEVEINCTRTQNTKCRCKPNFFCNSTVCEHCDPCTKCEHGIIKECTLTSNTKCKEEGS<br>RSNLGWLCLLLLPIPLIVWVKRKEVQKTCRKHRKENQGSHESPTLNPETVAINL GGSGGSG RIQNRPLVDLEHAALQ<br>VGKGIIPPPLREYGASEVRSVTRAFNHMAAGVKQLADDRTLLMAGVSHDLRTPLTRIRLATEMMSEQDGYLAESINKDIEECNAIIEQFIDYLR<br>TGQEMPMEMADLNAVLGEVIAAESGYEREIETALYPGSIEVKMHPLSIKRAVANMVVNAARYGNGWIKVSSGTEPNRAWFQVEDDGPGIAPEQR<br>KHLFQPFVRGDSARTISGTGLGLAIVQRIVDNHNGMLELGTSERGGLSIRAWLPVPVTRAQGTTKEG GGSG<br>VSKGEELFTGVVPILVELDGDVNGHKFSVSGEGEGDATYGKLTLKFICTTGKLPVPWPTLVTTFGYGLMCFARYPDHMKQHDFFKSAMPEGYVQ<br>ERTIFFKDDGNYKTRAEVKFEGDTLVNRIELKGIDFKEDGNILGHKLEYNYNSHNVYIMADKQKNGIKVNFKIRHNIEDGSVQLADHYQQNTPI<br>GDGPVLLPDNHYLSYQSALSKDPNEKRDHMVLLEFVTAAGITLGMDELYK DTYRYI<br><br>Brown: Fas receptor; Orange: EnvZ(C); Light green: mCitrine; Purple AU1 tag |
| **Constructs for input signal processing** | | |
| № | Name and scheme of construct | Amino acid or nucleotide sequence and description of parts |
| 50 | cTEV*-AP4-SbMVs-P3-nTEV<br>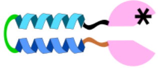 | M KSMSSMVSDTSCTFPSSDGIFWKHWIQTKDGQAGSPLVSTRDGFIVGIHSASNFTNTNNYFTSVPKNFMELLTNQEAQQWVSGWR<br>LNADSVLWGGHKVFMSKPEEPFQPVKEATQLMSELVYSQ YPYDVPDYA GGSGGGSGGS SPEDELAANEEELQQNEQKLAQIKQK<br>LQAIKYG GGSGGG ESVSLQS GSSGSG SPEDEIQQLEEEIAQLEQKNAALKEKNQALKYG EQKLISEEDL GESLFKGPRDYN<br>PISSTICHLTNESDGHTTSLYGIGFGPFIITNKHLFRRNNGTLLVQSLHGVFKVKNTTTLQQHLIDGRDMIIIRMPKDFPPFPQKLKFREPQRE<br>ERICLVTTNFQT<br><br>Magenta: cTEVp*; Green: AP4; Red: SbMVs cleavage site; Green: P3; Dark blue: Myc tag; Magenta: nTEVp |
| 51 | P4-PPVs-cTEV<br>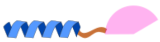 | M SPEDKIAQLKQKIQALKQENQQLEEENAALEY GGGSGGG NVVVHQA KSMSSMVSDTSCTFPSSDGIFWKHWIQTKDGQCGSP<br>LVSTRDGFIVGIHSASNFTNTNNYFTSVPKNFMELLTNQEAQQWVSGWRLNADSVLWGGHKVFMSKPEEPFQPVKEATQLMSELVYSQ<br>YPYDVPDYA<br><br>Green: P4; Cyan: PPVs cleavage site; Magenta: cTEVp; Dark blue: HA tag |
| 52 | SEAP-P4<br>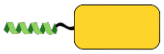 | M LLLLLLLGLRLQLSLGIIPVEEENPDFWNREAAEALGAAKKLQPAQTAAKNLIIFLGDGMGVSTVTAARILKGQKKDKLGPEIPL<br>AMDRFPYVALSKTYNVDKHVPDSGATATAYLCGVKGNFQTIGLSAAARFNQCNTTRGNEVISVMNRAKKAGKSVGVVTTTRVQHASPAGTYAHT<br>VNRNWYSDADVPASARQEGCQDIATQLISNMDIDVILGGGRKYMFRMGTPDPEYPDDYSQGGTRLDGKNLVQEWLAKRQGARYVWNRTELMQAS<br>LDPSVTHLMGLFEPGDMKYEIHRDSTLDPSLMEMTEAALRLLSRNPRGFFLFVEGGRIDHGHHESRAYRALTETIMFDDAIERAGQLTSEEDTL |

| | | |
|---|---|---|
| | | SLVTADHSHVFSFGGYPLRGSSIFGLAPGKARDRKAYTVLLYGNGPGYVLKDGARPDVTESESGSPEYRQQSAVPLDEETHAGEDVAVFARGPQ AHLVHGVQEQTFIAHVMAFAACLEPYTACDLAPPAGTTDAAHPGYSRVGAAGRFEQT<br>SGSG SPEDKIAQLKQKIQALKQENQQLEEENAALEYG<br><br>Magenta: SEAP; Green: P4 peptid |
| 53 | SEAP-P4-SbMVs-KDEL<br>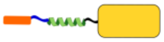 | M LLLLLLLGLRLQLSLGIIPVEEENPDFWNREAAEALGAAKKLQPAQTAAKNLIIFLGDGMGVSTVTAARILKGQKKDKLGPEIPL AMDRFPYVALSKTYNVDKHVPDSGATATAYLCGVKGNFQTIGLSAAARFNQCNTTRGNEVISVMNRAKKAGKSVGVVTTTRVQHASPAGTYAHT VNRNWYSDADVPASARQEGCQDIATQLISNMDIDVILGGGRKYMFRMGTPDPEYPDDYSQGGTRLDGKNLVQEWLAKRQGARYVWNRTELMQAS LDPSVTHLMGLFEPGDMKYEIHRDSTLDPSLMEMTEAALRLLSRNPRGFFLFVEGGRIDHGHHESRAYRALTETIMFDDAIERAGQLTSEEDTL SLVTADHSHVFSFGGYPLRGSSIFGLAPGKARDRKAYTVLLYGNGPGYVLKDGARPDVTESESGSPEYRQQSAVPLDEETHAGEDVAVFARGPQ AHLVHGVQEQTFIAHVMAFAACLEPYTACDLAPPAGTTDAAHPGYSRVGAAGRFEQT SGSG<br>SPEDKIAQLKQKIQALKQENQQLEEENAALEYG SGSG ESVSLQS KDEL<br><br>Magenta: SEAP; Green: P4 peptid; Black: SbMV protease cleavage site; Orange: KDEL signal |
| 54 | SS-P3-TEVs-KDEL<br>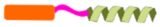 | M DMRVLAQLLGLLLLCFPGARC SPEDEIQQLEEEIAQLEQKNAALKEKNQALKYGS GSG ENLYFQS KDEL<br><br>Gray: Signaling sequence; Blue: P3 peptide; Black: TEV protease cleavage site; Orange: KDEL signal |

### Insulin secretion

| № | Name and scheme of construct | Amino acid or nucleotide sequence and description of parts |
|---|---|---|
| 55 | Preproinsulin_TEVs-KDEL<br>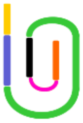 | M ALWMRLLPLLALLALWGPDPAAA FVNQHLCGSHLVEALYLVCGERGFFYTP RTKR EAEDLQVGQVELGGGPGAGSLQPLA LEGSL<br>RQKR GIVEQCCTSICSLYQLENYCN ENLYFQS KDEL<br><br>Gray: Signalling sequence; Cyan: Furin protease cleavage site; Green: C peptide Blue: B chain; Red: A chain; Black: TEV protease cleavage site; Orange: KDEL signal |
| 56 | Bchain_FURs_Gluc_FURS_Achain_TEVs_KDEL<br>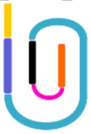 | M ALWMRLLPLLALLALWGPDPAAA FVNQHLCGSHLVEALYLVCGERGFFYTP RTKR KPTENNEDFNIVAVASNFATTDLDADR GKLPGKKLPLEVLKEMEANARKAGCTRGCLICLSHIKCTPKMKKFIPGRCHTYEGDKESAQGGIGEAIVDIPEIPGFKDLEPMEQFIAQVDLCV DCTTGCLKGLANVQCSDLLKKWLPQRCATFASKIQGQVDKIKGAGGD RQKR GIVEQCCTSICSLYQLENYCN ENLYFQS KDEL<br><br>Gray: Signalling sequence; Cyan: Furin protease cleavage site; Green: Gaussia luciferase; Blue: B chain; Red: A chain; Black: TEV protease cleavage site; Orange: KDEL signal |
| 57 | Preproinsulin<br>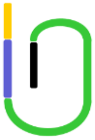 | M ALWMRLLPLLALLALWGPDPAAA FVNQHLCGSHLVEALYLVCGERGFFYTP RTKR EAEDLQVGQVELGGGPGAGSLQPLA LEGSL<br>RQKR GIVEQCCTSICSLYQLENYCN<br><br>Gray: Signalling sequence; Cyan: Furin protease cleavage site; Green: C peptide Blue: B chain; Red: A chain |

### FURp and KDELR1

| № | Name and scheme of construct | Amino acid or nucleotide sequence and description of parts |
|---|---|---|
| 58 | Furin protease<br>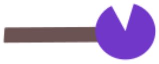 | M ELRPWLLWVVAATGTLVLLAADAQGQKVFTNTWAVRIPGGPAVANSVARKHGFLNLGQIFGDYYHFWHRGVTKRSLSPHRPRHSR LQREPQVQWLEQQVAKRRTKRDVYQEPTDPKFPQQWYLSGVTQRDLNVKAAWAQGYTGHGIVVSILDDGIEKNHPDLAGNYDPGASFDVNDQDP DPQPRYTQMNDNRHGTRCAGEVAAVANNGVCGVGVAYNARIGGVRMLDGEVTDAVEARSLGLNPNHIHIYSASWGPEDDGKTVDGPARLAEEAF FRGVSQGRGGLGSIFVWASGNGGREHDSCNCDGYTNSIYTLSISSATQFGNVPWYSEACSSTLATTYSSGNQNEKQIVTTDLRQKCTESHTGTS ASAPLAAGIIALTLEANKNLTWRDMQHLVVQTSKPAHLNANDWATNGVGRKVSHSYGYGLLDAGAMVALAQNWTTVAPQRKCIIDILTEPKDIG KRLEVRKTVTACLGEPNHITRLEHAQARLTLSYNRRGDLAIHLVSPMGTRSTLLAARPHDYSADGFNDWAFMTTHSWDEDPSGEWVLEIENTSE ANNYGTLTKFTLVLYGTAPEGLPVPPESSGCKTLTSSQACVVCEEGFSLHQKSCVQHCPPGFAPQVLDTHYSTENDVETIRASVCAPCHASCAT CQGPALTDCLSCPSHASLDPVEQTCSRQSQSSRESPPQQQPPRLPPEVEAGQRLRAGLLPSHLPEVVAGLSCAFIVLVFVTVFLVLQLRSGFSF RGVKVYTMDRGLISYKGLPPEAWQEECPSDSEEDEGRGERTAFIKDQSAL<br><br>Magenta: furin protease |
| 59 | KDELR1<br>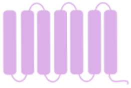 | M NLFRFLGDLSHLLAIILLLLKIWKSRSCAGISGKSQVLFAVVFTARYLDLFTNYISLYNTCMKVVYIACSFTTVWLIYSKFKATY DGNHDTFRVEFLVVPTAILAFLVNHDFTPLEILWTFSIYLESVAILPQLFMVSKTGEAETITSHYLFALGVYRTLYLFNWIWRYHFEGFFDLIA IVAGLVQTVLYCDFFYLYITKVLKGKKLSLPA<br><br>Purple: KDELR1 |

**Supplementary Table 2:** Constructs with a minimal promoter used in this study to measure the kinetics of secretion in systems based on the induction of transcription

| **Plasmid name and construct illustration** | **Plasmid description** |
|---|---|
| pcDNA 3.1/7x[b]_$P_{MIN}$_SEAP | Plasmid ensures expression of the SEAP reporter upon binding of a transcriptional activator to target sites upstream of a minimal promoter. The plasmid contains 7 copies of target sites [b]. |
| pcDNA 3.1/10[ab]_$P_{MIN}$_preproinsulin | Plasmid ensures expression of the preproinsulin upon binding of a transcriptional activator to target sites upstream of a minimal promoter. The plasmid contains 10 copies of target sites [ab]. |
| U6_gRNA[b] | Plasmid ensures constitutive expression of gRNA[b], driven from the murine U6 promoter. The gRNA targets a sequence designated [b]. |
| U6_gRNA[ab nt] | Plasmid ensures constitutive expression of gRNA[ab nt], driven from the murine U6 promoter. The gRNA targets the non-template (nt) strand of a sequence designated [ab]. |

**Supplementary Table 3**: Nucleotide sequence of the promoter regions used in this study

| **$P_{CMV}$** | CMV promoter from plasmid vector pcDNA3 |
|---|---|
| acattgattattgactagttattaatagtaatcaattacggggtcattagttcatagcccatatatggagttccgcgttacataacttacggtaaatggcccgcctg gctgaccgcccaacgacccccgcccattgacgtcaataatgacgtatgttcccatagtaacgccaatagggactttccattgacgtcaatgggtggactatttacgg taaactgcccacttggcagtacatcaagtgtatcatatgccaagtacgccccctattgacgtcaatgacggtaaatggcccgcctggcattatgcccagtacatgac cttatgggactttcctacttggcagtacatctacgtattagtcatcgctattaccatggtgatgcggttttggcagtacatcaatgggcgtggatagcggtttgact cacggggatttccaagtctccaccccattgacgtcaatgggagtttgttttggcaccaaaatcaacgggactttccaaaatgtcgtaacaactccgccccattgacg caaatgggcggtaggcgtgtacggtgggaggtctatataagcagagctc | |
| **$P_{MIN}$** | Minimal promoter |
| tagagggtatataatggaagctcgacttccag | |
| **U6 promoter** | |
| gatccgacgcgccatctctaggcccgcgccggccccctcgcacagacttgtgggagaagctc ggctactcccctgccccggttaatttgcatataatatttcctagtaactatagaggcttaatgtgcgataaaagacagataatctgttcttt ttaatactagctacattttacatgataggcttggatttctataacttcgtatagcatacattatacgaagttataaacagcacaaaaggaaa ctcaccctaactgtaaagtaattgtgtgttttgagactataagtatcccttggagaaccaccttgttgg | |
| **7x gRNA binding site [b]** | tcttccgtttccacatct |
| ccgtcttccgtttccacatcttcacggataccaaggtggcaccggtttgattgcacagcccgtcttccgtttccacatcttcacggtgctggcagccggggtaccgcaggca agtcgcccgtcttccgtttccacatcttcacggatggaatcacggcggccgcccgtatctcatccccgtcttccgtttccacatcttcacggaatgcgtcggcgtgcggccg ctgccagttgcaccgtcttccgtttccacatcttcacggttggctgtcctagggtacctggacgctccttgccgtcttccgtttccacatcttcacggtggctggtgccgcacc ggtaagcagacggccgccgtcttccgtttccacatcttcacggggatcctgtacgggccaga | |
| **10x gRNA binding site[ab]** | gcttcttccgtttccacatct |
| Cggtggcaccggtttgattgcacagctttactgctgctcccgcttcttccgtttccacatcttgctggcagccggggtaccgcaggcaagtcgctttactgctgctcccgctt cttccgtttccacatctatggaatcacggcggccgcccgtatctcatcctttactgctgctcccgcttcttccgtttccacatctaatgcgtcggcgtgcggccgctgccagttg catttactgctgctcccgcttcttccgtttccacatctttggctgtcctagggtacctggacgctccttgtttactgctgctcccgcttcttccgtttccacatcttggctggtgccg caccggtaagcagaactagaggtggcaccggtttgattgcacagctttactgctgctcccgcttcttccgtttccacatcttgctggcagccggggtaccgcaggcaagtc gctttactgctgctcccgcttcttccgtttccacatctatggaatcacggcggccgcccgtatctcatcctttactgctgctcccgcttcttccgtttccacatctaatgcgtcggc gtgcggccgctgccagttgcatttactgctgctcccgcttcttccgtttccacatctttggctgtcctagggtacctggacgctccttgtttactgctgctcccgcttcttccgtttc cacatct | |

**Supplementary Table 4.** The amount of transfected plasmids in each well of 96-well plate used for realization of individual logic functions using the SEAP activity readout

**Microscopy, Figure 1c, Supplementary figure 1a**

| Input plasmid | ng | Transfection control | ng |
|---|---|---|---|
| TagRFP_FURS_TD_TEVs_KKYL | 70 | tagBFP | 50 |
| TagRFP_FURs_TD_TEVs_KKMP | 70 | tagBFP | 50 |
| TagRFP_TEVs_KDEL | 70 | tagBFP | 50 |
| TagRFP_TD_TEVs | 70 | tagBFP | 50 |

**Westerm blot, Supplementary figure 1 b, c**

| Input plasmid | ng |
|---|---|
| SEAP-TEVs-KDEL | 1500 |
| SEAP-3xFURs-TD-TEVs-KKYL | 1500 |

**membER and lumER secretion, Figure 1d, e, Supplementary figure 2**

| Input protease | ng | Reporter plasmid | ng |
|---|---|---|---|
| erTEVP | 0<br>10 | SEAP_TEVs_KDEL | 50 |
| erSbMVp | 0<br>10 | SEAP_SbMVs_KDEL | 50 |
| TEVp | 0<br>80 | SEAP_3xFURs_TM_3xTEVs_KKYL | 60 |
| PPVp | 0<br>80 | SEAP_3xFURs_TM_3xPPVs_KKYL | 60 |
| SbMVp | 0<br>80 | SEAP_3xFURs_TM_3xSbMVs_KKYL | 60 |
| SuMMVp | 0<br>80 | SEAP_3xFURs_TM_3xSuMMVs_KKYL | 60 |
| TEVp | 0<br>80 | SEAP_3xFURs_TM_3xTEVs_KKMP | 60 |
| TEVp | 0<br>20<br>40<br>80 | SEAP_3xFURs_TM_3xTEVs_KKMP | 60 |
| erTEVp | 0<br>0.4<br>1.2<br>5<br>10 | SEAP_3xFURs_TM_3xTEVs_KKYL | 60 |
| erTEVp | 0<br>10<br>20<br>40<br>80 | Gluc _TEVs_KDEL | 60 |
| TEVp | 0<br>60 | SEAP_3xFURs_TM_3xTEVs_KKYL | 60 |
| FURp | 0<br>10<br>20<br>40<br>80 | | |
| erPPVp | 0<br>5<br>10<br>20 | SEAP_PPVs_KDEL | 50 |
| erPPVp (N23Q, T173G) | 0<br>5<br>10<br>20 | SEAP_PPVs_KDEL | 50 |

**membER and lumER inducible secretion, Figure 2b, Supplementary figure 3, 4**

| Input plasmids | ng | Reporter plasmid | ng | Inducer molecule |
|---|---|---|---|---|
| FKBP_cerTEVp:<br>FRB_nerTEVp | 20:20<br>40:40<br>60:60 | SEAP_TEVs_KDEL | 90 | rapamycin |
| FKBP_cTEVp:<br>FRB_nTEVp | 20:20<br>40:40<br>60:60 | SEAP_3xFURs_TM_3xTEVs_KKYL | 90 | rapamycin |
| ABI_cTEVp:<br>PYL1_nTEVp | 20:20<br>40:40<br>60:60 | SEAP_3xFURs_TM_3xTEVs_KKYL | 90 | ABA |
| FKBP_cTEVp:<br>FRB_nTEVp | 20:20<br>40:40<br>60:60 | SEAP_3xFURs_TM_3xTEVs_KKMP | 90 | rapamycin |
| FKBP_cerTEVp:<br>FRB_nerTEVp | 20:20<br>40:40<br>60:60<br>80:80 | SEAP_TEVs_KDEL | 20<br>40<br>60<br>80 | rapamycin |
| ABI_cerTEVp<br>PYL1_nerTEVp | 10:10<br>20:20<br>40:40 | SEAP_TEVs_KDEL | 20<br>40<br>60<br>80 | ABA |

**Secretion kinetics, Figure 2c, d, Supplementary figure 5, 6**

| Input plasmids | ng | Reporter plasmid | ng | Inducer molecule |
|---|---|---|---|---|
| FKBP_cerTEVp:<br>FRB_nerTEVp | 20:20 | SEAP_TEVs_KDEL | 60 | rapamycin |
| FKBP_cTEVp:<br>FRB_nTEVp | 80:80 | SEAP_3xFURs_TM_3xTEVs_KKYL | 60 | rapamycin |
| ABI_cTEVp:<br>PYL1_nTEVp | 80:80 | SEAP_3xFURs_TM_3xTEVs_KKYL | 60 | ABA |
| dCas9:FKBP<br>FRB:VPR<br>sgRNA [b] | 25<br>25<br>25 | pMIN_SEAP | 60 | rapamycin |
| dCas9:ABI<br>PYL1:VPR<br>sgRNA [b] | 25<br>25<br>25 | pMIN_SEAP | 60 | ABA |

**Addition of KDELR and FURp, Supplementary figure 2f, 7**

| Input plasmids | ng | Reporter plasmid | ng |
|---|---|---|---|
| TEVp<br>FURp | | SEAP_3xFURs_TM_3xTEVs_KKYL | 1000 |
| erTEVp<br>KDELR | | SEAP_TEVs_KDEL | 1000 |

**Secretion with added Eeyarestatin I, Supplementary figure 8**

| Input plasmids | ng | Reporter plasmid | ng | Inducer molecule |
|---|---|---|---|---|
| FKBP_cerTEVp<br>FRB_nerTEVp | 150<br>150 | SEAP_TEVs_KDEL | 1000 | rapamycin |
| FKBP_cTEVp<br>FRB_nTEVp<br>FURp | 600<br>600<br>100 | SEAP_3xFURs_TM_3xTEVs_KKYL | 1000 | rapamycin |

**Microscopy – lumER secretion, Figure 3a,b, Supplementary figure 9**

| Input plasmid | ng | Protease | ng | Transfection control | ng |
|---|---|---|---|---|---|
| GFP11x7-Fas-Env<br>SS-GFP1-10-TEVs-KDEL | 150<br>150 | erTEVp | 30 | iRFP | 20 |
| GFP11x7-Fas-Env<br>SS-GFP1-10-TEVs-KDEL | 150<br>150 | FKBP_cerTEVp:<br>FRB_nerTEVp | 30<br>30 | iRFP | 20 |

**Microscopy – membER secretion, Figure 3c,d, Supplementary figure 10**

| Input plasmid | ng | Protease | ng | Transfectio n control | ng |
|---|---|---|---|---|---|
| SS-GFP1-10-FURs-KD-TEVs-KKYL<br>GFP11x7-Fas-Env-iRFP | 100<br>100 | FKBP_cTEVp:<br>FRB_nTEVp | 100<br>100 | iRFP | 20 |
| SS-GFP1-10-FURs-KD-TEVs-KKYL<br>GFP11x7-Fas-Env-iRFP | 100<br>100 | TEVp | 100<br>100 | iRFP | 20 |

| membER and lumER orthogonality, Figure 4 | | | | |
|---|---|---|---|---|
| **Input plasmids** | **ng** | **Reporter plasmid** | **ng** | **Inducer molecule** |
| erTEVp | 10 | Bchain_Myc_Gluc_Achain_TEVs_KDEL : SEAP_FURs_TD_TEVs_KKYL | 15 30 | / |
| TEVp | 80 | | | |
| FKBP_cerTEVp : FRB_nerTEVp | 10:10 | Bchain_Myc_Gluc_Achain_TEVs_KDEL : SEAP_FURs_TD_TEVs_KKYL | 15 30 | ABA/rapamycin |
| ABI_cTEVp: PYL1_nTEVp | 80:80 | | | |

| Logical processing, Figure 5, Supplementary figure 12 | | | |
|---|---|---|---|
| **Input plasmids** | **ng** | **Reporter plasmid** | **ng** |
| cTEV*-AP4-SbMVs-P3-nTEV | 10 | SEAP_3xFURs_TD_TEVs_KKYL | 30 |
| P4-PPVs-cTEV | 5 | | |
| PPVp | 90 | | |
| SbMVp | 60 | | |
| cTEV*-AP4-SbMVs-P3-nTEV | 10 | SEAP_FURs_TD_TEVs_KKYL | 30 |
| P4-PPVs-cTEV | 5 | | |
| SbMVp | 60 | | |
| PPVp | 40 | | |
| P3-TEVs-KDEL | 20 | SEAP-P4 | 10 |
| erTEVp | 5 | | |
| erSbMVp | 5 | SEAP-P4-SbMVs-KDEL | 10 |
| P3-TEVs-KDEL | 40 | SEAP-P4-SbMVs-KDEL | 10 |
| erTEVP | 5 | | |
| erSbMVp | 5 | | |

| OFF switch, Figure 6, | | | | | |
|---|---|---|---|---|---|
| **Input plasmid** | **ng** | **Protease** | **ng** | **Protease** | **ng** |
| SEAP_3xFURs_TD_TEVs_KKYL | 20 | FKBP_PPVs_cTEVp<br>FRB_PPVs_nTEVp | 30<br>30 | PPVp | 0<br>20<br>40<br>80<br>120 |
| | | TEVp | 30 | | |
| SEAP_3xFURs_TD_TEVs_KKYL<br>furin | 300<br>100 | FKBP_PPVs_cTEVp<br>FRB_PPVs_nTEVp | 300<br>300 | ABI_cPPVp<br>PYL_nPPVp | 800<br>800 |

| Insulin secretion, Figure 7, Supplementary figure 13 | | | | |
|---|---|---|---|---|
| **Input plasmids** | **ng** | **Input plasmids** | **ng** | **Inducer molecule** |
| FKBP_cerTEVp:<br>FRB_nerTEVp | 70<br>70 | Preproinsulin_TEVs_KDEL | 550 | rapamycin |
| dCas9:ABI | 150 | pMIN_preproinsulin | 450 | ABA |
| PYL1:VPR | 150 | | | |
| sgRNA | 150 | | | |
| erTEVp | 0<br>10<br>20<br>50<br>100 | Bchain_Gluc_Achain_TEVs_KDEL | 20 | / |
| FKBP_cerTEVp:<br>FRB_nerTEVp | 0:0<br>5:5<br>10:10<br>20:20<br>30:30 | Bchain_Gluc_Achain_TEVs_KDEL | 20 | rapamycin |

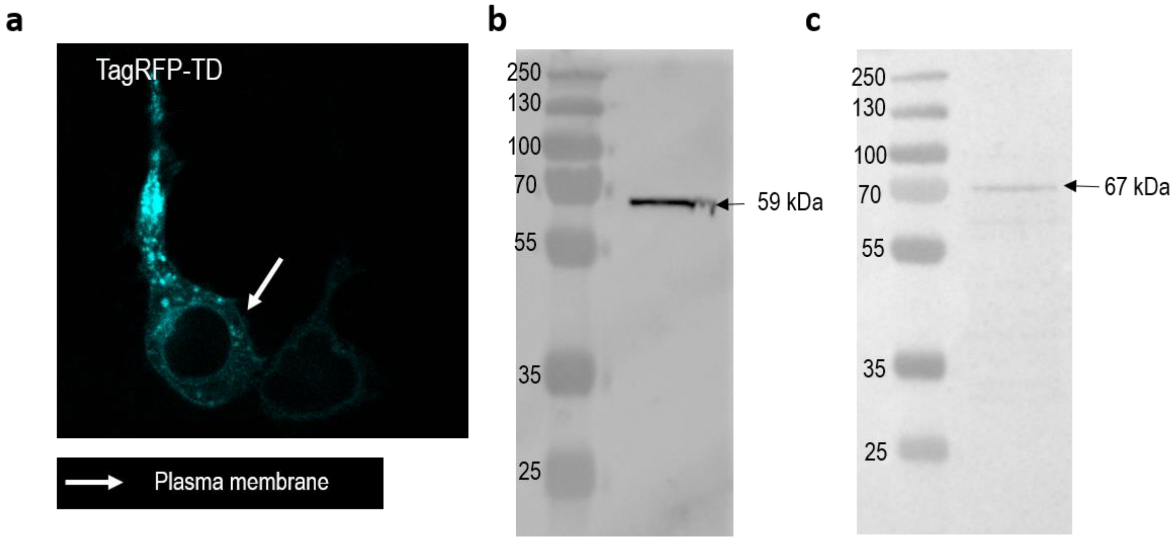


**Supplementary figure 1: Expression pattern of lumER and membER constructs. a,** TagRFP-TD-TEVs, lacking a C-terminal retention signal, was visualized in HEK293T cells. In contrast to constructs featuring a retention sequence (Figure 1b-d), TagRFP signal is detected also at the plasma membrane in the absence of a retention signal, indicating that the protein trafficked to the cell surface. Plasmids coding either SEAP-TEVs-KDEL (AU1 tag) (**b**) or SEAP-FURs-TD-TEVs-KKYL (AU1 tag) (**c**) were transfected into HEK293T, lysed after 48h and their expression was verified by Western blot. A band at 59 kDa corresponds to SEAP-TEVs-KDEL and a band at 67 kDa corresponds to SEAP-FURs-TD-TEVs-KKYL.

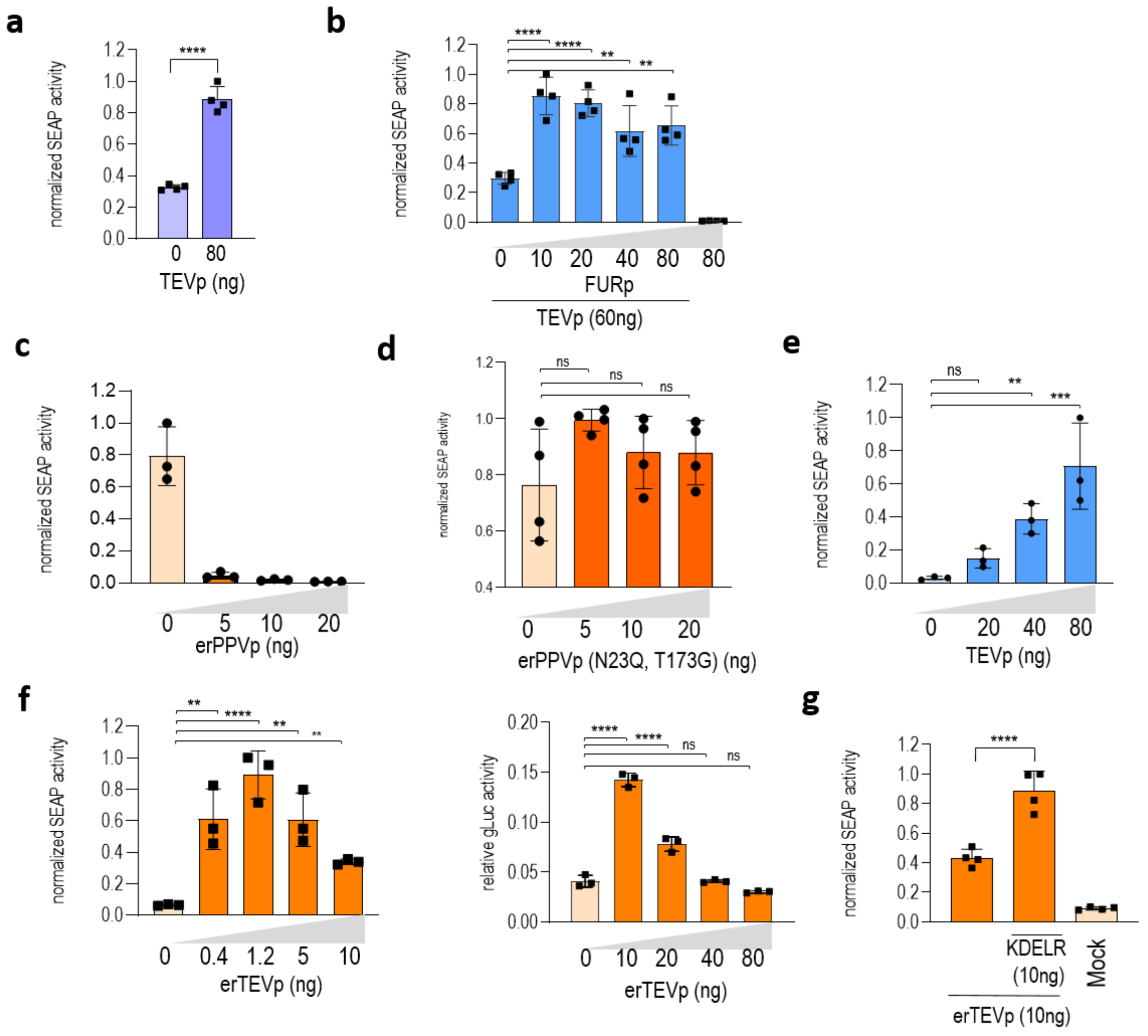


**Supplementary figure 2: Proteolysis-based protein secretion.** **a**, Effect of a TEVp-cleavable C-terminal KKMP ER retention sequence. **b**, Co-transfection of furin protease (FURp) increases the amount of secreted SEAP in the membER system. **c**, Regulation of protein secretion by the erPPVp targeting SEAP-PPVs-KDEL. The protease lacks the catalytic activity inside ER. **d**, Regulation of protein secretion of a mutated version of erPPVp(N23Q, T173G) targeting SEAP-PPVs-KDEL. **e**, TEVp concentration dependence on protein secretion with the membER system, detected by the secreted SEAP. **f**, erTEVp concentration dependence on protein secretion with the lumER system, detected by the secreted SEAP (left) and Gaussia luciferase (Gluc, right). **f**, Cotransfection of KDELR1 increases the retention capacity of the system. Values are the mean of three or four cell cultures ± s.d and are representative of two independent experiments. Significance was tested by a one-way analysis of variance (ANOVA) with Dunnett's comparison.

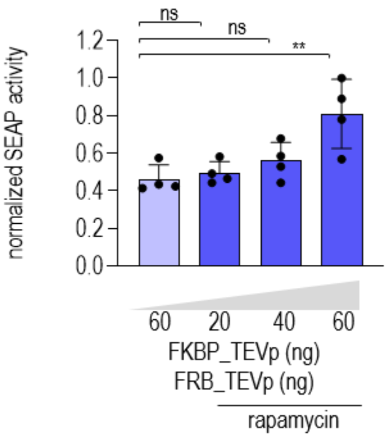


**Supplementary figure 3: Rapamycin inducible secretion of the membER system with a KKMP retention signal.** Values are the mean of four cell cultures ± s.d. and are representative of two independent experiments. Significance was tested by a one-way analysis of variance (ANOVA) with Dunnett's comparison.

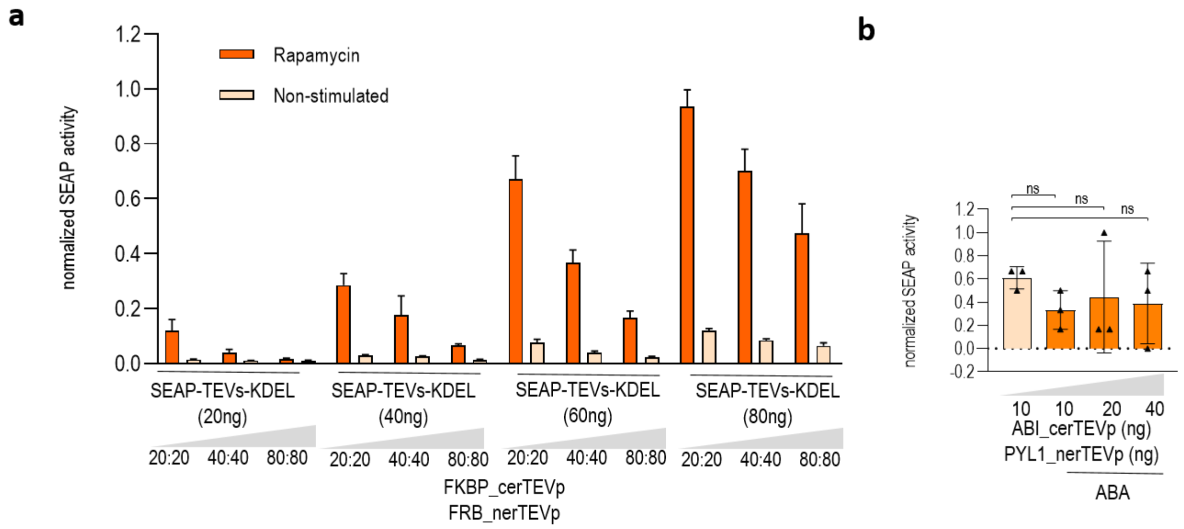


**Supplementary figure 4: Inducible secretion of SEAP by CID with the lumER system. a,** 20, 40 or 80 ng of pcDNA3.1 comprising $P_{CMV}$_FKBP-cerTEVp and $P_{CMV}$_FRB-nerTEVp (er-rapa-TEV) were co-transfected with 20, 40, 60 or 90ng of pcDNA $P_{CMV}$_SEAP-TEVs-KDEL. SEAP activity in the medium was measured 16h after stimulation with rapamycin. Increasing the amount of the split protease decreases the fold difference in secreted SEAP between stimulated and non-stimulated cells. **b**, Regulation of proteins secretion by ABI-cerTEV/PYL1-nerTEVp and SEAP-TEVs-KDEL. The dimerization potential of ABI/PYL1 domains is diminished inside of ER (er-aba-TEV). Values are the mean of three or four cell cultures ± s.d. and are representative of two independent experiments. Significance was tested by a one-way analysis of variance (ANOVA) with Dunnett's comparison.

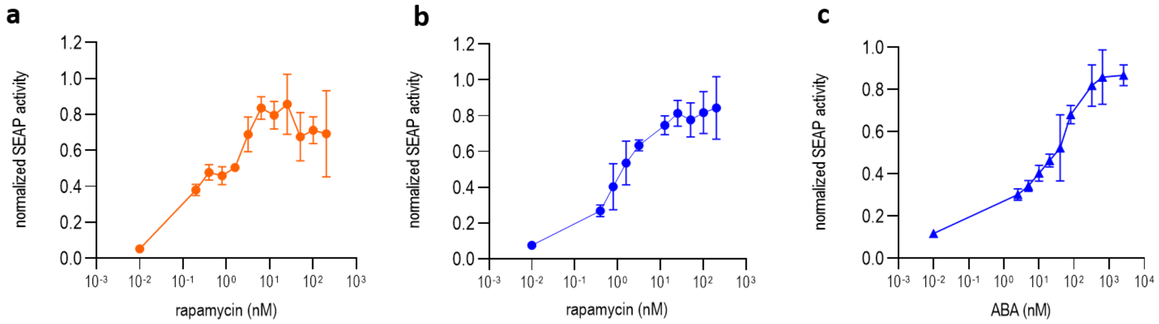


**Supplementary figure 5: SEAP secretion dependence on the concentration of chemical inducer. a,** Titration of rapamycin on SEAP-TEVs-KDEL (lumER system). **b**, Titration of rapamycin on SEAP-FURs-TM-TEVs-KKYL (membER system). **c**, Titration of ABA on SEAP-FURs-TM-TEVs-KKYL (membER system). Values are the mean of three cell cultures ± s.d. and are representative of two independent experiments.

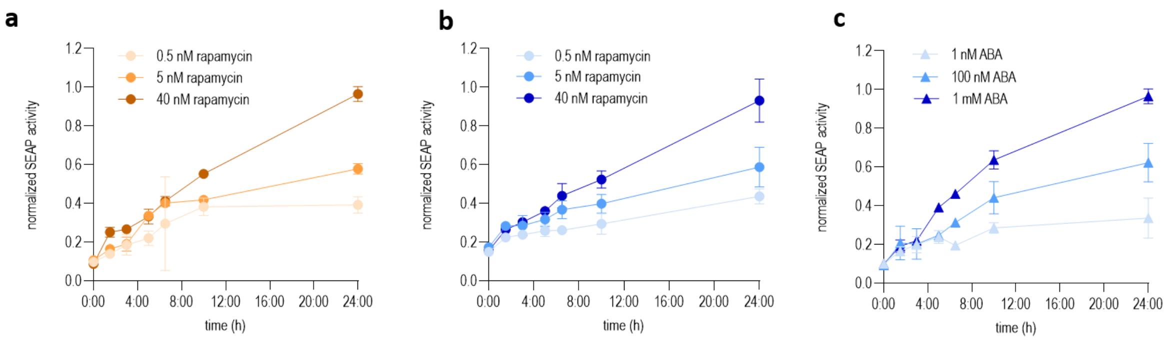


**Supplementary figure 6: Secretion kinetics of lumER and membER with varying concentration of chemical inducer.** HEK293T cells were transfected with either the lumER (orange, panel **a**) or membER (blue, panels **b** and **c**) system in combination with an inducible split TEVp. Two days after transfection cells were stimulated with rapamycin (round symbols) at a final concentration of 0.5 nM, 5nM and 40 nM, or ABA (triangle symbols) at a final concentration of 1 nM, 100 nM and 1 mM. Media was sampled at specific time points and SEAP activity was measured. Values are the mean of three cell cultures ± s.d. and are representative of two independent experiments.

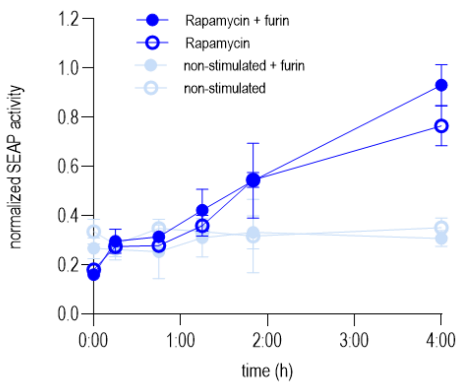


**Supplementary figure 7: The effect of furin protease on the kinetics of SEAP secretion with the membER system.** Co-transfection of FURp did not significantly impact the secretion kinetics in the first 4h after induction of secretion. Values are the mean of four cell cultures ± s.d and are representative of two independent experiments.

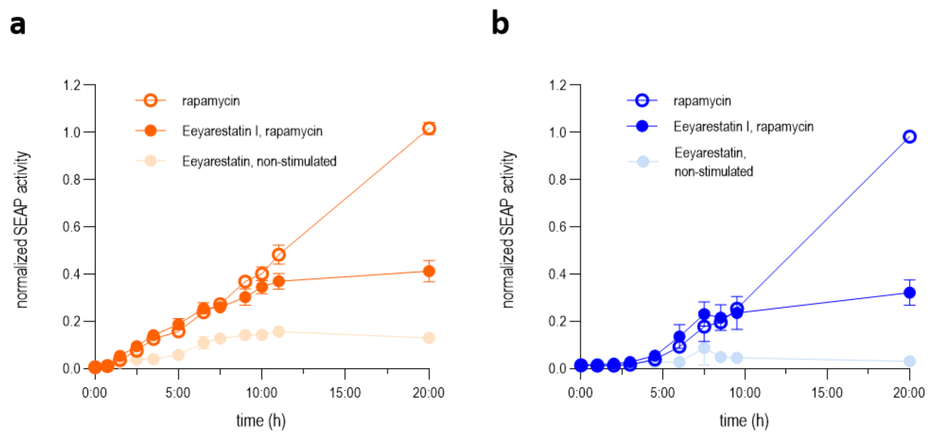


**Supplementary figure 8: Retention capacity of secretion systems.** Cells were treated with an inhibitor of ER transport, Eeyarestatin I. After stimulation with rapamycin, cell medium was harvested at specified time points and SEAP activity was measured. Both the lumER (**a**) and membER (**b**) system release most of the stored protein around 10h after induction of secretion. The remaining secretion in cells not treated with Eeyarestatin I is most likely due to the re-storing of ER with newly synthesized protein. Values are the mean of three cell cultures ± s.d. and are representative of two independent experiments.

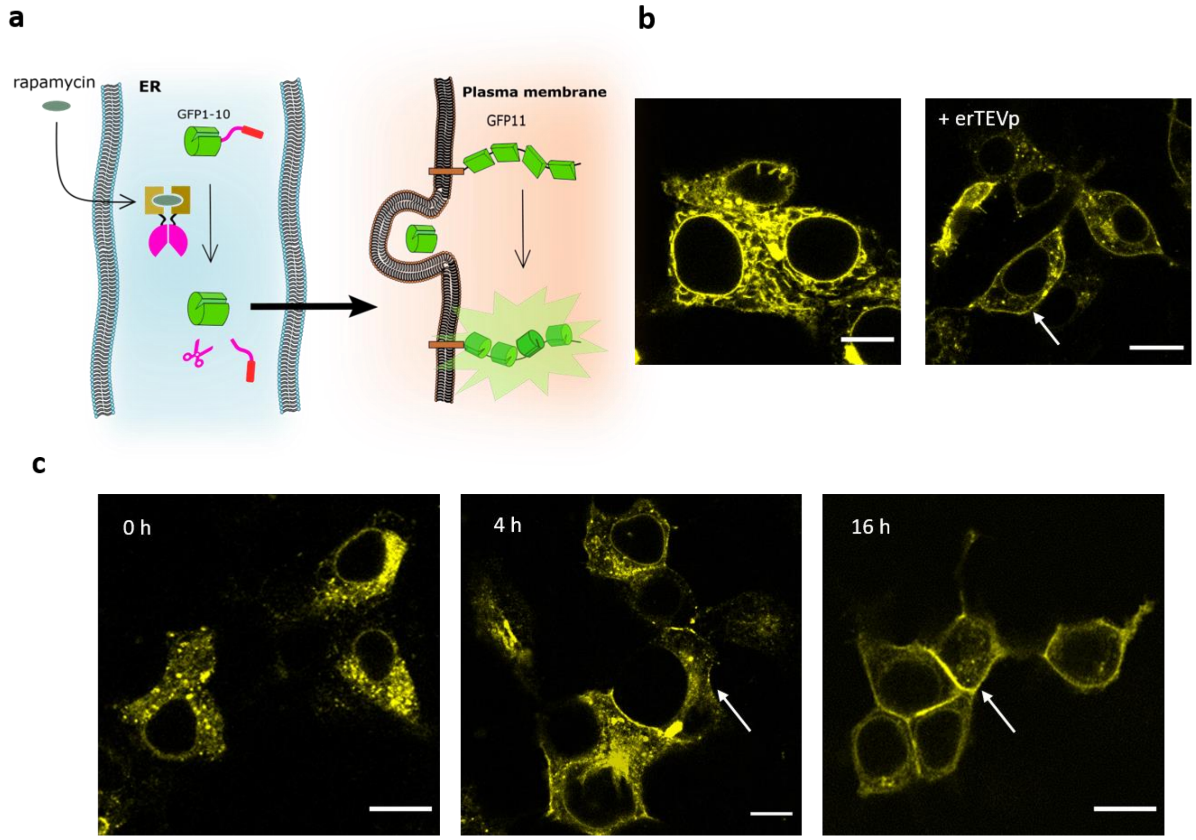


**Supplementary figure 9: Visualization of protein trafficking in the lumER system.** A split GFP fragment, composed of the 10 β-strands of the fluorescent protein (GFP1-10), was fused to the lumER system, while the complementary fragment, composed of 7 repeats of the 11$^{th}$ β-strand (GFP11x7), was fused to the FAS receptor transmembrane domain, localizing it to the plasma membrane. Removal of the retention signal allows GFP1-10 to move to the membrane and associate with its complementary part, reconstituting full GFP, which can be detected under the microscope. **b**, GFP fluorescence was measured before and after co-transfection of erTEVp. After the addition of the protease, the signal was detected on the cell membrane (white arrow), where it was previously undetected. **c**, FKBP-cerTEVp /FRB-nerTEVp was used inducible reconstitution of the split protease and the removal of the retention signal. GFP fluorescence was visualized before stimulation of cells with rapamycin, 4h after and 16h after the addition of rapamycin. Scale bar = 10 μm

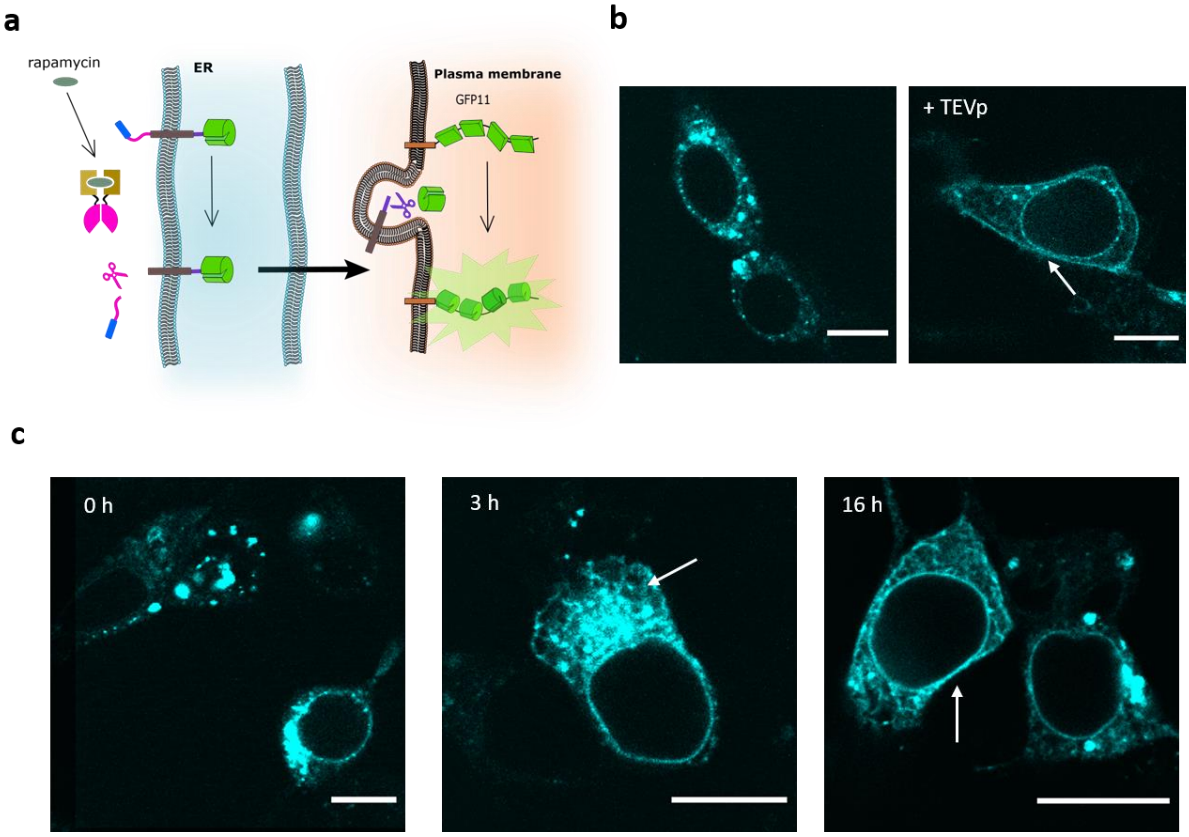


**Supplementary figure 10: Visualization of protein trafficking in the membER system.** Similar as the lumER system, a GFP1-10 was fused to the membER system, while the complementary fragment, GFP11x7, was fused to the FAS receptor transmembrane domain. A cytosolic protease facilitates the removal of the KKYL signal, allowing GFP1-10 to move to its complementary fragment, thus regaining its fluorescent activity. **b**, GFP fluorescence was measured before and after co-transfection of TEVp. After the addition of the protease, the signal was detected on the cell membrane (white arrow), where it was previously undetected. **c**, FKBP-cTEVp /FRB-nTEVp was used inducible reconstitution of the split protease and the removal of the retention signal. GFP fluorescence was visualized before stimulation of cells with rapamycin, 3h after and 16h after the addition of rapamycin. Scale bar = 10 µm

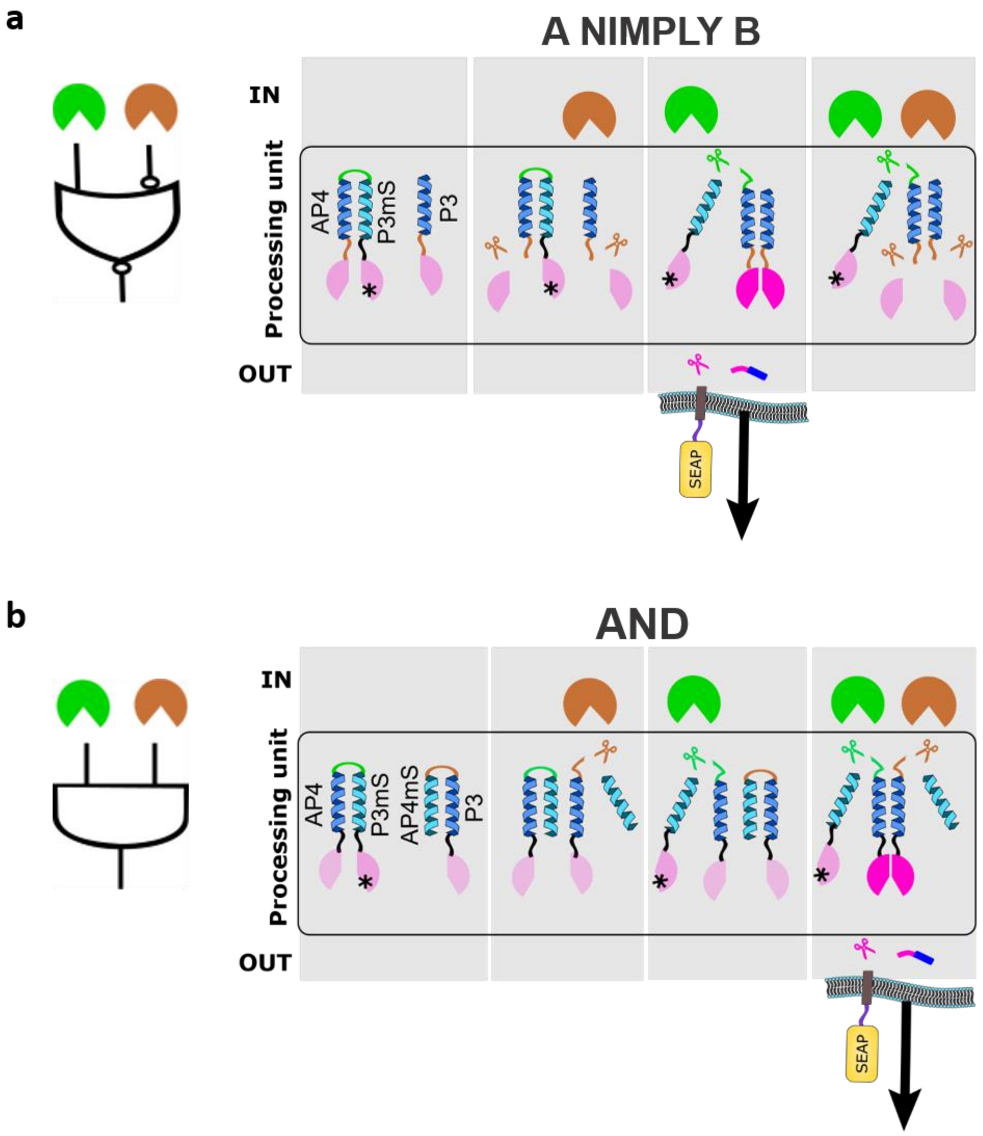


**Supplementary figure 11: Schematic representation of the constructs used for A nimply B and AND Boolean SPOC logic circuits used to regulate membER-SEAP secretion . a**, design of A nimply B logical function. The constructs used for signal processing were nTEV-SbMVs-AP4-PPVs-P3mS-cTEV* and P3-SbMVs-cTEV. An output in the form of a reconstituted TEVp, which further regulates the secretion of SEAP, is produced only when PPVp is present, but not SbMVp. This is achieved through the introduction of an inhibitory coil P3mS, which prevents the reconstitution of the two protease fragments through the interaction of the complementary P3/AP4 coiled-coils. An introduction of an inactive cTEV fragment (indicated with cTEV*) further prevents unwanted coupling of the split protease fragments. The removal of the inhibitory P3mS-cTEV* is achieved by the cleaving of the PPVs (green line), which is positioned in the linker connecting the AP4/P3mS coiled coils. A negative regulation with SbMVp is achieved by positioning a SbMVs (brown line) between the active fragments of the split TEVp and its adjacent coil. **b**, design of AND logical function. The constructs used for AND signal processing were nTEV-AP4-PPVs-P3mS-cTEV* and AP4mS-SbMVs-P3-nTEV. In this setting both PPVp and SbMVp need to be present in order to remove the P3mS-cTEV* and AP4mS inhibitory coils and allow for the coupling of the split TEVp fragments through P3/AP4 coiled-coil interaction.

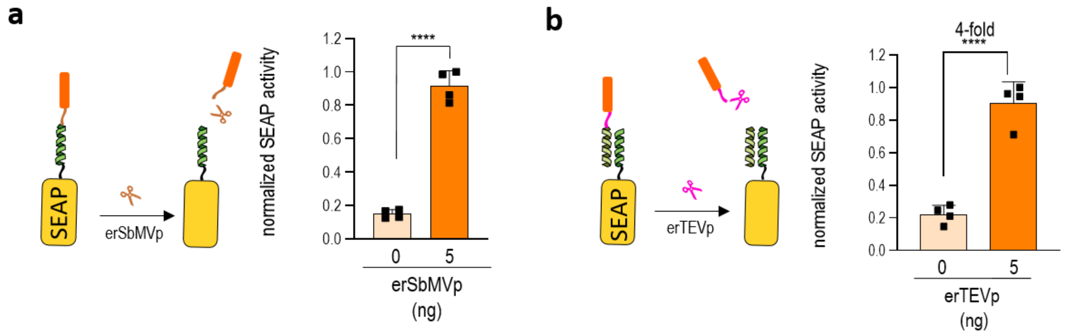


**Supplementary figure 12: Retention of SEAP inside the ER through coiled-coil interactions. a**, SEAP was fused to a P4 coiled-coil forming peptide and a KDEL retention signal, separated by SbMVs. Upon the addition of erSbMVp, secretion is induced. **b**, Addition of P3-TEVs-KDEL significantly decreases the secretion of SEAP-P4, which itself does not contain a retention signal. Upon the addition of erTEVp, SEAP is detected in the media. Values are the mean of four cell cultures ± s.d. and are representative of two independent experiments. An unpaired two-tailed t test (after equal variance was assessed with the F test assuming normal data distribution) was used for the statistical comparison of the data.

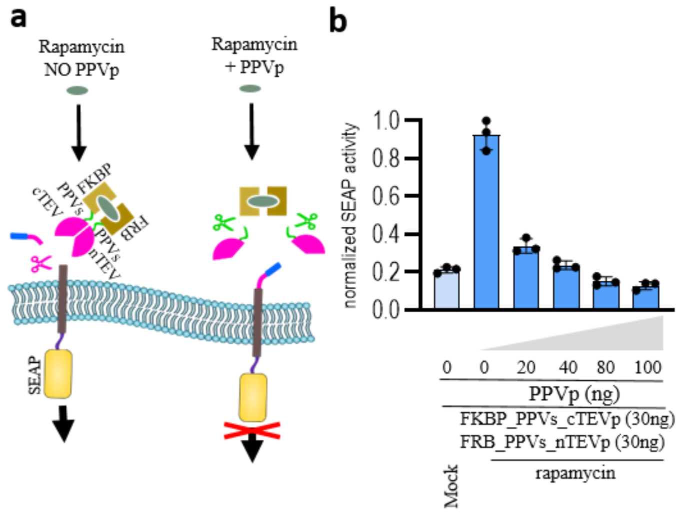


**Supplementary figure 13: A simplified A nimply B logical function for the membER system. a,** A PPVp cleavage site (PPVs) was inserted between the FKBP/FRB dimerization domains and TEV protease fragments. The reconstitution of TEVp can be inhibited by the addition of PPVp, **b**, Cells were stimulated with rapamycin to induce secretion. Cells co-expressing PPVp showed decreased secretion of SEAP, compared to cells where no PPVp was added. Values are the mean of three cell cultures ± s.d. and are representative of two independent experiments.

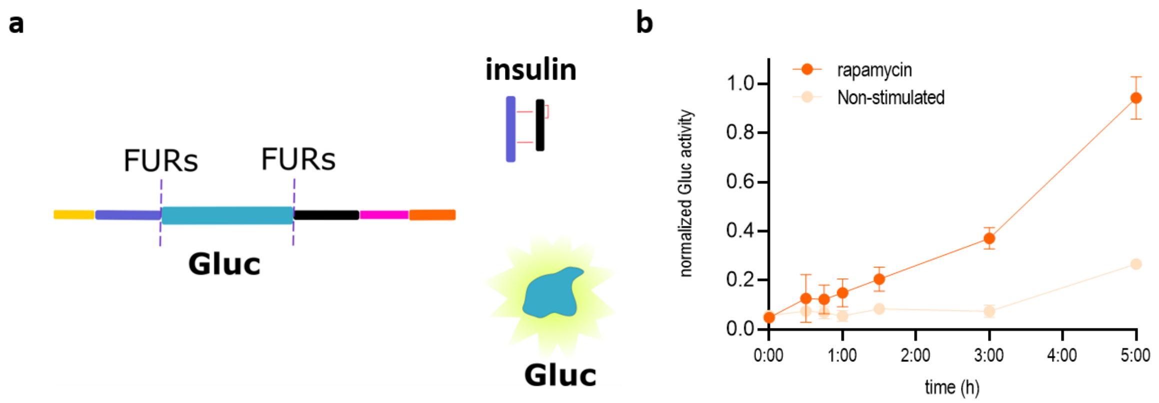


**Supplementary figure 14: Monitoring insulin secretion by measuring Gluc activity in the media.** As an alternative system to monitor the release of insulin into the cell media after the addition of rapamycin, the C-peptide sequence was replaced by Gaussia luciferase (Gluc) and luciferase activity was measured in the media as a proxy for insulin secretion. Values are the mean of four cell cultures ± s.d. and are representative of two independent experiments.
